# Flexible working memory through selective gating and attentional tagging

**DOI:** 10.1101/846675

**Authors:** Wouter Kruijne, Sander M. Bohte, Pieter R. Roelfsema, Christian N. L. Olivers

**Affiliations:** Faculty of Behavior and Movement Sciences, Vrije Universiteit Amsterdam, Amsterdam, Noord Holland, The Netherlands; Machine Learning Group, Centrum voor Wiskunde & Informatica, Amsterdam, Noord Holland, The Netherlands; Department of Vision & Cognition, Netherlands Institute for Neuroscience, Amsterdam, Noord Holland, The Netherlands; Department of Integrative Neurophysiology, Center for Neurogenomics and Cognitive Research, Vrije Universiteit Amsterdam, Amsterdam, Noord Holland, The Netherlands; Psychiatry Department, Academic Medical Center, Amsterdam, Noord Holland, The Netherlands

## Abstract

Working memory is essential for intelligent behavior as it serves to guide behavior of humans and nonhuman primates when task-relevant stimuli are no longer present to the senses. Moreover, complex tasks often require that multiple working memory representations can be flexibly and independently maintained, prioritized, and updated according to changing task demands. Thus far, neural network models of working memory have been unable to offer an integrative account of how such control mechanisms are implemented in the brain and how they can be acquired in a biologically plausible manner. Here, we present WorkMATe, a neural network architecture that models cognitive control over working memory content and learns the appropriate control operations needed to solve complex working memory tasks. Key components of the model include a gated memory circuit that is controlled by internal actions, encoding sensory information through untrained connections, and a neural circuit that matches sensory inputs to memory content. The network is trained by means of a biologically plausible reinforcement learning rule that relies on attentional feedback and reward prediction errors to guide synaptic updates. We demonstrate that the model successfully acquires policies to solve classical working memory tasks, such as delayed match-to-sample and delayed pro-saccade/antisaccade tasks. In addition, the model solves much more complex tasks including the hierarchical 12-AX task or the ABAB ordered recognition task, which both demand an agent to independently store and updated multiple items separately in memory. Furthermore, the control strategies that the model acquires for these tasks subsequently generalize to new task contexts with novel stimuli. As such, WorkMATe provides a new solution for the neural implementation of flexible memory control.

**Author Summary:** Working Memory, the ability to briefly store sensory information and use it to guide behavior, is a cornerstone of intelligent behavior. Existing neural network models of Working Memory typically focus on how information is stored and maintained in the brain, but do not address how memory content is controlled: how the brain can selectively store only stimuli that are relevant for a task, or how different stimuli can be maintained in parallel, and subsequently replaced or updated independently according to task demands. The models that do implement control mechanisms are typically not trained in a biologically plausible manner, and do not explain how the brain learns such control. Here, we present WorkMATe, a neural network architecture that implements flexible cognitive control and learns to apply these control mechanisms using a biologically plausible reinforcement learning method. We demonstrate that the model acquires control policies to solve a range of both simple and more complex tasks. Moreover, the acquired control policies generalize to new situations, as with human cognition. This way, WorkMATe provides new insights into the neural organization of Working Memory beyond mere storage and retrieval.

## Introduction

Complex behavior requires flexible memory mechanisms for dealing with information that is no longer present to the senses, but that is still relevant to current task goals. For example, when on the highway, before we decide it is safe to change lanes, we sequentially accumulate evidence in memory from various mirrors and the road ahead of us. Importantly, such complex behavior requires memory operations beyond mere storage: Not every object that we observe on the highway needs to be memorized, while often it is a specific combination of information (e.g. multiple cars and signs) which determines whether it is safe to switch lanes. As any novice driver has experienced, learning to properly apply these operations of selecting, maintaining, and managing the correct information in memory can take quite some effort. Yet, after sufficient practice, we learn to apply these skills and abstract the essence across multiple environments, regardless of the specifics of the road or cars around us. This example illustrates the core functions that define *Working Memory* (WM), and that – in the words of [1] – make WM *work*. First, Working Memory is *flexible* in that control processes determine what information is stored, when it is updated, and how it is applied during task performance. Second, the rules that govern these control operations for a given task setting are *trainable* and can be acquired with practice. Third, after training, these rules then *generalize* to the same task setting with different stimuli. It is this combination of flexibility, trainability, and generalizability that makes WM a cornerstone of cognition, not only in humans, but also in nonhuman primates [2–4]. Here we present a neural network model of WM that integrates these core components.

Before we describe the WorkMATe model, we will briefly explain how it extends previous models that either focused on the generic storage of arbitrary sensory stimuli in memory or on the learning of content-specific memory operations.

## Models of storage and matching

A number of previous neural network models explain how the brain can temporarily maintain information [5–9] and how different items can be maintained separately [10–13]. Given their emphasis on storage, one of the most commonly modeled WM tasks is the delayed match-to-sample task, in which the observer responds according to whether an observed stimulus matches a memorized stimulus.

Delayed matching tasks do not require an agent to act on the specific *content* of information in memory. Rather, the agent produces a response based on the presence or absence of sufficient *similarity* between two successively presented stimuli – which in principle could be anything. Experimental work has revealed that both human and nonhuman primates can almost effortlessly determine such matches, even for stimuli that have never been seen before [2, 3, 14, 15], and studies have demonstrated neurons in frontal as well as parietal cortices whose activity depends on the match between sensory input and memory content [16–18]. Taken together,these findings suggest that the computations governing matching tasks – determining the similarity or degree of match – is relatively independent of stimulus content.

Most models for matching, recognition, and recall tasks implement a content-independent computation of a match signal. [19] have demonstrated that a relatively simple, self-organizing neural network can learn to detect co-activation neuronal pools representing similar information with non-overlapping codes, allowing for a match signal between sensory and memory information to emerge upon presentation. Match signals also emerge in models of associative memory that assume one-shot Hebbian learning of arbitrary information in the hippocampus. In these models, the ease of subsequent context-driven retrieval provides an index of stimulus-memory similarity, which is used to simulate recall probabilities and response times [20–24]. [25] have shown that repetition suppression in the inferotemporal cortex after a repeated presentation of a stimulus predicts recognition performance for arbitrary stimuli in the macaque (see also [26, 27]). The same idea is prevalent in models of visual search, where a match signal is computed between an item in memory and stimuli present in the to-be-searched scene, which is subsequently used to optimally guide attention [28–30].

In these models, match signals are automatically computed as an emergent consequence of the interaction between perception and memory, adding to the utility of WM without the need for training on specific stimulus content first. In contrast, more complex tasks call for additional WM operations, decisions, and motor actions depending on specific content. In the lane changing example, an empty rear view mirror may indicate that overtaking is safe, unless the side mirror says otherwise. Generic match models typically do not explain how memory content is controlled, how control policies can be acquired through training, and how memory content in combination with sensory information leads to action selection. Such trainable, flexible, action-oriented models of WM will be discussed next.

## Models of memory operations

A rather different class of models has focused on how WM can be trained to solve tasks in which multiple different stimuli map onto different responses – that is, how the cognitive system learns which of a number of available actions, including memory operations, is appropriate given particular (combinations of) stimuli. Training neural network models to solve tasks means that as the network processes examples, weights are updated to establish a desirable mapping between an input and output stream. In a reinforcement learning setting, a desired, optimal mapping yields a policy that maximizes reward and minimizes punishment. For multilayer neural networks, this becomes a problem of ‘structural credit assignment’, where the learning algorithm needs to determine to what extent a connection weight contributed to the outcome. For memory tasks, there is an additional ‘temporal credit assignment’ problem, as the outcome of certain actions – e.g., storing an item into memory – will only later in the trial lead to success or failure. An ongoing issue in deep learning is how these credit assignment problems might be solved in a biologically plausible manner [31–35].

One biologically plausible solution to temporal and structural credit assignment in WM tasks is provided by the AuGMEnT algorithm [36–38], which in turn is based on the AGREL model for perceptual learning [39–41]. These models demonstrate that *attentional feedback* can play a critical role in solving credit assignment [42]. The architecture used by AuGMEnT is a multilayer neural network with a recurrent memory layer to maintain information. The output of the neural network is the expected reward value associated with each action. Upon selection of an action, the attentional feedback mechanism ‘tags’ synapses that contributed to this action. When an action does not yield the expected reward, a Reward Prediction Error (RPE) signal is broadcast across the network, which drives weight changes in tagged synapses. Through these mechanisms, AuGMEnT implements a rudimentary but trainable WM architecture.

This architecture can learn to solve a variety of memory tasks where sequences of stimuli need to be integrated over time to yield a correct response [36, 37]. However, at the same time, AuGMEnT lacks the operations that define the flexibility of primate WM: its store accumulates relevant information, but does not allow, for example, items to be separately updated, selectively forgotten, or only to be encoded under certain conditions.

A highly popular neural network architecture that does incorporate such flexible control mechanisms is the Long-Short-Term Memory (LSTM) architecture [43]. This architecture introduces a *gated* memory store, implemented through gating units that open or close dependent on activity in the rest of the network. These gates allows an agent to control which information is allowed entry into memory, how new information is integrated, and which information is read out at any given time. LSTM networks and similar architectures are now commonplace in modern deep learning systems, which is a testament to their power [44–50]. However, while LSTM architectures allow for flexible control over memory content, they were not developed with biological plausibility in mind: typical implementations rely on rather implausible learning rules from a biological perspective [43, 49]. LSTMs can be trained using reinforcement learning methods [51, 52], but the complexity of the recurrent architecture renders training implausibly inefficient when compared to animal learning (requiring millions of trials to learn a relatively straightforward T-maze task).

One of the best established biologically inspired models of flexible WM control so far, is the Prefrontal Cortex-Basal Ganglia Working Memory model (PBWM [1, 53, 54]). PBWM allows for flexible memory control in a manner inspired by LSTM, but was designed with a strong focus on biologically plausibility. PBWM only gates the entry of sensory stimuli into its WM store in an all-or-none fashion. Specifically, the basal ganglia determine whether items are allowed to enter WM, on the basis of selecting internal gating actions. The model can learn complex hierarchical tasks (such as 12-AX, described below), which require selective updating and maintenance of relevant items in WM while preventing the storage of distractor stimuli. However, as noted by [55], the exact functionality of PBWM is somewhat obscured by the fact that it is a rather complex model with a highly interwoven architecture of neural subsystems and several parallel learning algorithms, both supervised and unsupervised [56–58]. [55] demonstrated that a much simpler reinforcement learning architecture, that distills only PBWMs gating mechanism, could learn the same type of tasks. Thus, these trainable, action-oriented models (AuGMEnT, LSTM and PBWM) illustrate how neural networks can learn tasks that go beyond mere storage and matching. In both LSTM and PBWM, memory control is flexible: items can be encoded, maintained and updated separately, and there are mechanisms that prevent interference from task-irrelevant stimuli. Notably, and in stark contrast with more storage-oriented models, these models solve tasks by constructing memory representations tailored to the task at hand: sensory information is encoded in a manner that makes them fit to solve the task, and that link them to relevant actions. In a more conceptual wording: these networks store *task-relevant content*, rather than generic copies of sensory stimuli. They provide control operations to update specific content and learn to apply them based on reinforcement. Yet, these models do not easily cope with arbitrary stimuli that the agent has never observed before. For this, the models lack the generic storage approach that matching-oriented models utilize, and it remains untested whether generalized matching signals can be integrated in this type of model.

## WorkMATe: generalizable, flexible, trainable WM

As laid out above, existing neural network models of flexible memory vary according to their focus of functionality (storage vs. action). Here, we present WorkMATe (Working Memory through Attentional Tagging) a neural network architecture that integrates the core components of these models to arrive at a biologically plausible model of WM that is trainable, flexible and generalizable. The model utilizes a new, gated memory circuit inspired by both PBWM and LSTM, to maintain multiple items separately in WM. We include a straightforward neuronal circuit for a generic matching process that compares the memory content to incoming new stimuli. Last but not least, these structures are embedded in a multilayer neural network that is trained using the simple and biologically plausible reinforcement learning rule of AuGMEnT. We demonstrate how the resulting neural network architecture solves complex, hierarchical tasks with multiple stimuli that have different roles depending on context, and that it can rapidly generalize an acquired task policy to novel stimuli that it has never encountered before.

## Materials and methods

We will first describe the architecture of WorkMATe and how it compares the memory representations to sensory stimuli, as well as how its biologically plausible learning rule resolves the credit assignment problem by combining reinforcement learning with an attentional feedback mechanism. We will then illustrate the virtues of WorkMATe in four general versions of popular WM tasks. First, we model a basic delayed match-to-sample task with changing stimulus sets to illustrate how the model generalizes to novel stimuli. Second, we illustrate hierarchical problem solving with the classic “memory juggling” 12-AX task, where the agent is presented with a stream of symbols and must learn the rule. Third, the challenges of both these tasks are combined by training it on a sequential recognition task introduced by [2, 3], where an agent has to store multiple, sequentially presented items, and match them to subsequent test stimuli, in the same order. Here again we assess both flexibility and generalization to new stimuli. Finally, we turn to the delayed pro-saccade/anti-saccade task [59–63], because it allows for a direct comparison between the present architecture and the previous ‘gateless’ AuGMEnT model [36].

We present a model architecture (Fig. 1A) that achieves good performance in these four memory tasks. The parameter values and other network specifics were kept the same in all simulations. An overview of these parameters is given in supporting table S1 Table. The details of these computations will be described here, followed by a discussion of our simulations. Code used to implement the architecture and run the simulations is available from an online repository, accessible via https://osf.io/jrkdq/?view_only=e2251230b9bf415a9da837ecba3a7d64.

**Fig 1.**
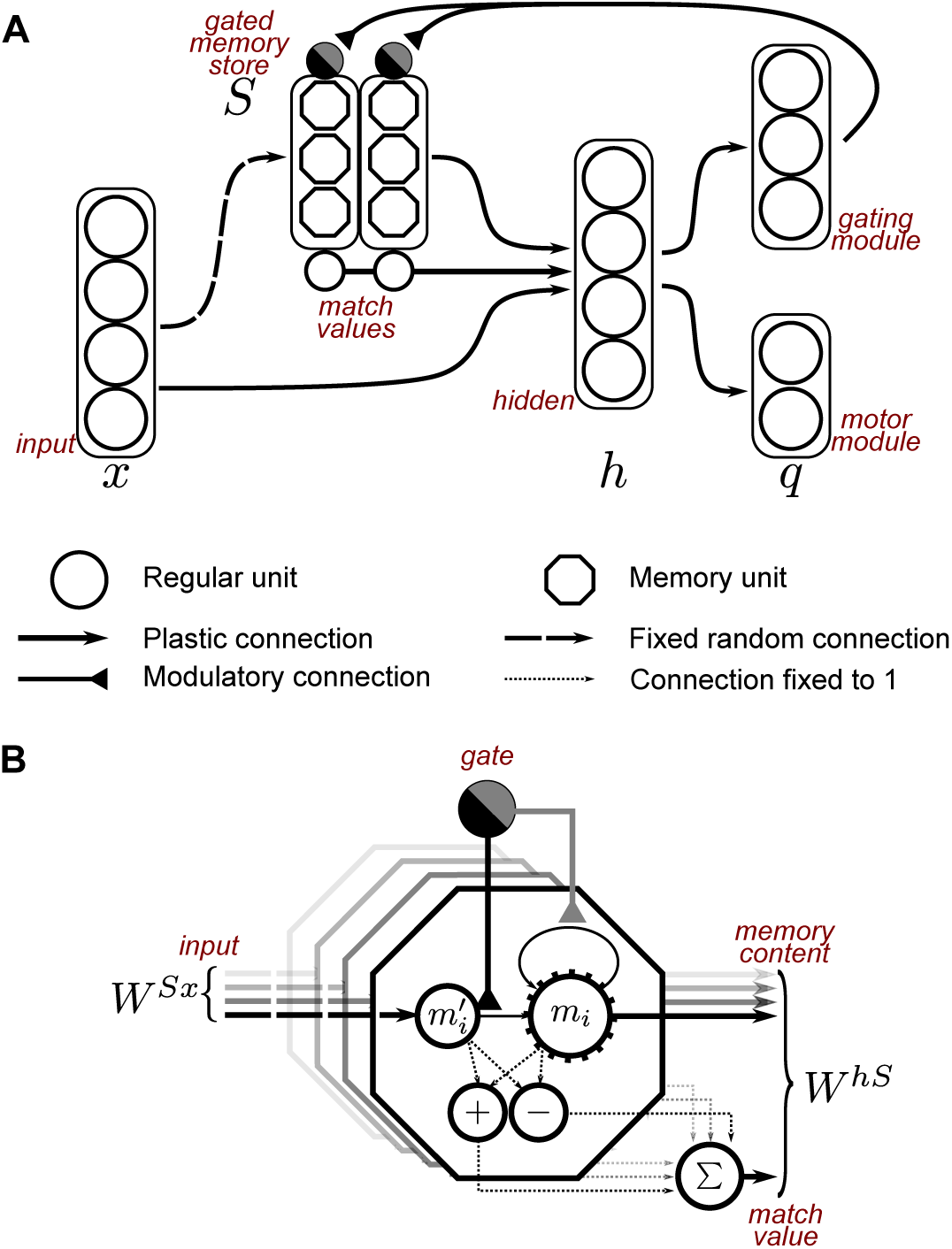
A The network architecture used in all simulations: a standard multilayer network, complemented by a gated store composed of two independent memory blocks. The input layer and memory store both project to the hidden layer, which in turn projects to two output modules. There, activity encodes Q-values that drive action selection. **B** Memory unit within a block, with a ‘closed’ gate: the memory content is maintained via self-recurrent connections. Additionally, a match value is computed between sensory and memory information, by comparing a projection of the sensory information (*m*′_*i*_) to memory content (*m*_*i*_). The comparison is performed by two units which respond to positive and negative disparities between the two values. Their output is summed across memory units, yielding one match value for each block. The closed gate inhibits the connection between *m*′_i_ → *m*_*i*_ so that the original memory is maintained. Only when a gating action is selected, the recurrent projection is inhibited and *m*′_*i*_ → *m*_*i*_ is opened so that memory content is updated. Fig. 2 illustrates network activity in a task context, and supporting table S1 Table lists the number of units in each layer.

### Input representations and Feed-forward sweep

The model is a neural network that receives input *x* at every time step *t*. Input is composed of sensory representations *x*_*s*_ and a representation of time *x_τ_*. Sensory representations are, in all simulations, defined binary patterns with activity levels [1, 0] that uniquely identify each stimulus. The time representation is inspired by ‘*time* cells’ as identified in multiple cortical and sub-cortical areas [4, 23, 64, 65]. These cells encode time by their delayed response profiles, each peaking at different times relative to the onset of a trial. In our model units, activity profiles are defined as symmetrically increasing and decreasing activity around each unit’s unique peak time, at the levels [0.125, 0.25, 0.5, 1.0]. An example sequence of inputs from Time-and Sensory units is depicted in Fig. 2C.

**Fig 2.**
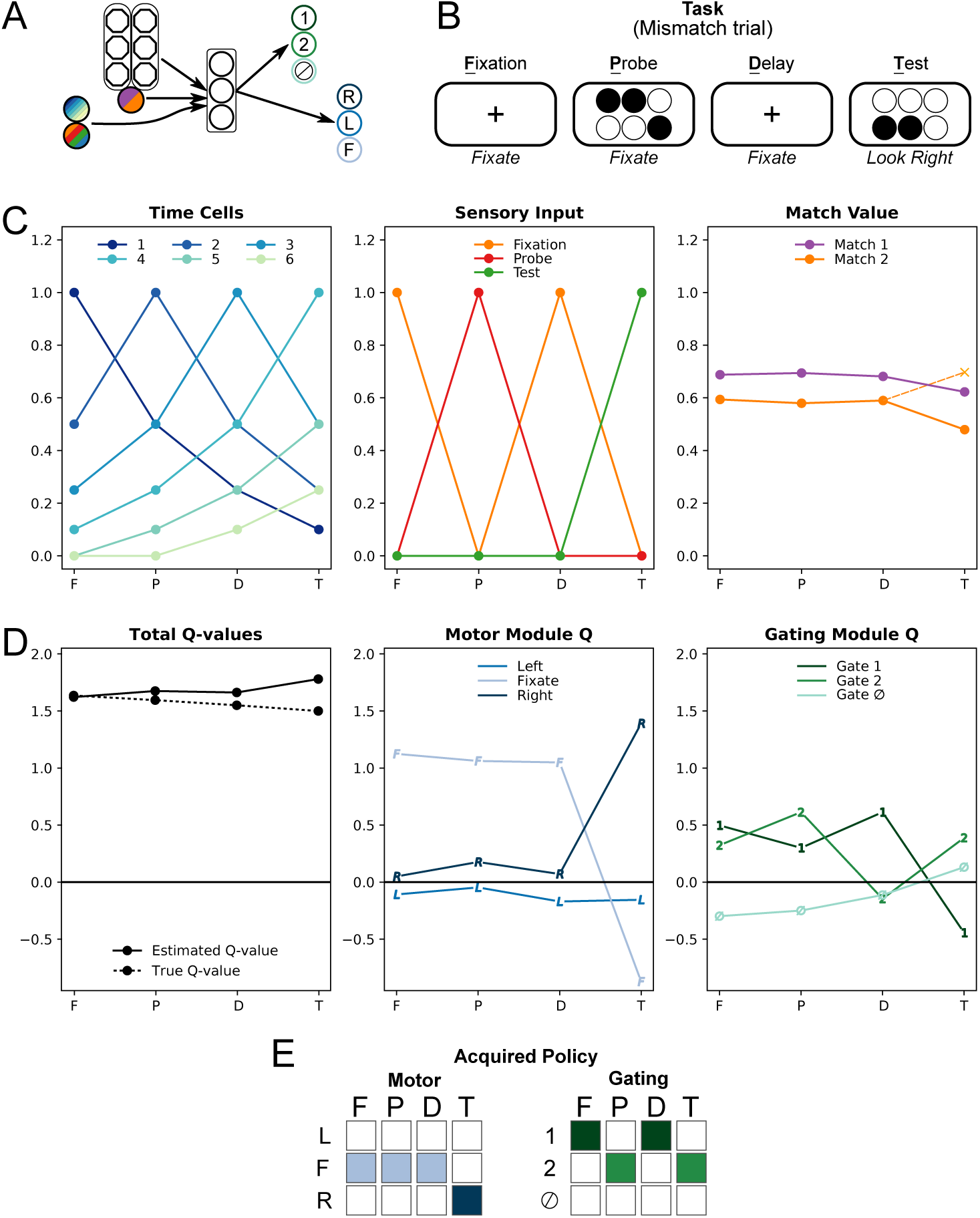
Neural Network performance on an illustrative ‘mismatch’ trial, after training. **A:** Model architecture (as in Fig. 1). Colors in the input-and output layers correspond to the graphs in **C** and **D**. **B:** Example mismatch trial. After fixation, a probe stimulus (a 6-bit pattern) is presented which must be maintained in memory. After a delay, the test stimulus is presented. The agent must hold fixation, and then indicate whether the test matches the probe stimulus (leftwards saccade) or not (rightwards saccade) **C:** From left to right: Time units *x_τ_* (6 out of 10 inputs shown), sensory units *x*_*s*_, and match values *x*_*m*_ at different time steps during the trial (F-P-D-T as in **B**, see main text for details). **D:** After training, the agent estimates the Q-value of its selected action set (left) from the Q-values in the motor-and gating module (middle and right). **E:** Motor-and gating policy, i.e. vector *z* at each time point determined through winner-take-all selection over the *q*-values in **D.** The colored squares indicate the chosen internal and external actions at each time step. Colors correspond to those in **D**.

The network projects the input representation *x* to two different layers. One is a regular hidden layer *h* in which units are activated via the projection weight matrix *W^hx^*. The other layer, activated through projection *W^Sx^* is the memory store *S*, which is composed of two equally sized memory blocks *m*_1_ and *m*_2_. During the initial feedforward sweep of activity, the projection *W*^*Sx*^⋅ *x* = *S*′ = {*m*′_1_, *m*′_2_} serves to compute the match value between the projected sensory representation *m^l^_i_*and the contents of each memory block *m*_*i*_. These match values are denoted as *x*_*m*1_ *, x_m_*_2_. As noted above, there are a number of hypotheses on how this match value might be computed [19, 20, 24, 25]. Here, we refrained from a specific modeling effort and instead computed the sum of absolute differences between the two representations, which is a common match metric [66–68]:

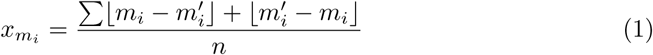

Here, *n* refers to the number of nodes in a block. This computation could be readily implemented by a set of accessory units that are activated by input projections and memory units, and respond to disparities between memory-and sensory information (Fig. 1B). The summed activation in these units is a measure of dissimilarity, and one minus this value is used as a match signal.

The current activity of the memory circuit *S^∗^* = {*S, x_m_*}, which reflects the memory content as well as the match values, is projected to the hidden layer *h*. This regular hidden layer integrates information from the input layer and memory stores, via:

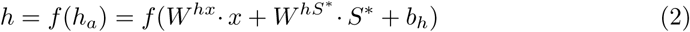

where *b* is a bias input vector, and *f* is a standard sigmoid transfer function.

The hidden layer *h* projects to the output layer *q* through the weights *W^qh^*:

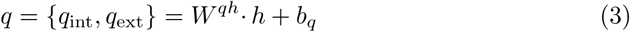

these output values *q* will, after training, approximate the Q-values of each of the possible response options. The Q-value is the sum of the expected immediate and temporally discounted future rewards for the remainder of the trial that the agent can acquire by selecting that action. The output layer *q* is divided into two modules: one for *external* and one for *internal* actions. External actions reflect the motor response options of the agent, which in our simulations reflect either holding or releasing a lever, or fixating left, right, or in the center of the screen. The internal actions determine memory gating. Based on the action selected in this layer, the currently presented stimulus is either memorized in block 1, in block 2, or it is ignored. On most time steps, the agent will selects internal and external actions associated with the units with the highest activation {**argmax**(*q*_int_), **argmax**(*q*_ext_)}. On rare exploration time steps, (determined by exploration rate *ϵ*), the agent will select a random action, determined by a Boltzmann controller operating over the q-values within each module.

### Storage and Gating

The memory layer *S* in WorkMATe is functionally similar to that used in PBWM. Separate memory representations are maintained via self-recurrent projections in the memory store. This is a strong abstraction of the neurophysiological mechanisms of WM maintenance in the primate brain, as there is no consensus in the literature as to whether items in WM are functionally organized into slots [69, 70], continuous resources [71–73], hierarchically organized feature bundles [74], or through interactions with long-term memory representations [75]. Here, we remain largely agnostic regarding the precise representation, but choose a mechanism where items in memory can be maintained separately, can be updated separately, and can be selectively ignored to prevent interference [1]. We will show that this approach allows us to investigate how complex cognitive control over the content of WM can be acquired via reinforcement learning.

After feedforward processing is completed and the Q-values in the output layer have been computed, the agent selects a gating action from {*g*_1_, *g*_2_,*g*_∅_}, in order to either gate the current sensory representation into block *m*_1_*, m*_2_ or to prevent the stimulus from entering the memory store altogether. Note that unlike in PBWM, a memory representation *m*_*i*_ is not a direct copy of sensory information. Rather, it is a compressed representation of the input representation, encoded via the weights *W^Sx^*. This allows for generalization of learned task rules to novel stimuli.

Importantly, unlike the other, trained projections in the model, *W^Sx^* remains fixed throughout each model run at the connection strengths it obtains through random initialization. As a result, memory representations of a stimulus are not tuned to the task at hand, and will differ dependent on whether they are encoded in block 1 or block 2. Previous work [76–78] has demonstrated that utrained random projections can be used for memory encoding in a useful manner, as long as dissociable memory representations can be formed. This is not to say that memory encoding in the brain is necessarily random and untrained, but we will use this architecture to illustrate that without additional tuning, the model can successfully encode stimuli in a generic manner, an will explore whether learned policies generalize to novel stimulus sets.

### Learning

Learning in the model follows the AuGMEnT-algorithm [36], which was in turn derived from the AGREL learning rule [39]. At every time step, the model predicts the Q-value of each of its possible actions. These values are represented in the motor-and the gating-module in the network’s output layer. Based on these values, the gating module selects an internal action and the motor module an external action, in parallel. The sum of the two Q-values associated with the selected actions, *q*_int_(*t*) + *q*_ext_(*t*), reflect the total Q-value *Q*_*t*_, i.e. the network’s estimate of the sum of discounted rewards predicted for the remainder of the trial. Note that there is no a priori constraint on how these two values are weighted, though in all the tasks simulated here we found the Q-values in internal and external action modules to converge to comparable values, with each module accounting for approximately half of the total Q-value associated with the selected pair of actions.

The selected actions form a binary vector *z*, which is 1 for the units reflecting the selected actions, and 0 otherwise. AuGMEnT/AGREL states that once actions have been selected, an attentional feedback signal originates from these units which passes through the system through attentional feedback connection. This recurrent signal is used to ‘tag’ synapses that contributed to the selected actions. These synaptic tags correspond to eligibility traces in traditional *SARSA*(*λ*) reinforcement learning. The value of these tags gradually decays at each time step with a rate *α* = 1 − *λγ*, where *γ* is a temporal discounting factor. (discussed below). The update of a tag depends on the contributions of a synapse to a selected action. Formally, this means that in each plastic connection in the weight matrices *W^Sx^, W^hx^, W^hS∗^, W^qh^*, each Tag_*ji*_ between presynaptic unit *i* and postsynaptic unit *j* is updated according to:

for the connections *h →* q:

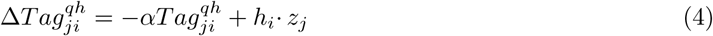

and for the connections *x → h* and *S → h*:

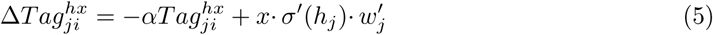

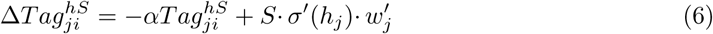

with:

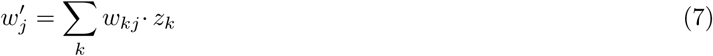

Here, the term *h*_*j*_ refers to the output of hidden unit *j*, and *σ*′ is the derivative of the sigmoid transfer function. The term *w_j_^l^* indicates the amount of recurrent feedback from the action vector *z* onto the hidden layer nodes. This feedback is determined by the weight between the hidden nodes and the selected actions where *z*_*k*_ = 1 if action k is selected, and *z*_*j*_ = 0 for all non-selected actions *j* Feedback connections are updated via the same learning rule as the feedforward connections. Therefore, the feedforward and feedback connections remain or become reciprocal, which has been observed in neurophysiology [79].

Synaptic connections are updated when the synaptic tags interact with a global reward-prediction error (RPE) signal. This signal, *δ*(*t*) is modeled after striatal dopamine, and reflects the signed difference between the expected and obtained reward. This is expressed in the SARSA temporal difference rule:

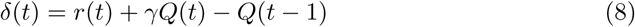

That is, the model values the previous actions on the basis of the obtained reward *r*(*t*) plus the amount of expected future reward *Q*(*t*) multiplied by a temporal discounting factor *γ* ϵ [0, 1], and contrasts this valuation with the previously expected value *Q*(*t* − 1). The RPE then triggers a global, neuromodulatory signal that spreads uniformly throughout the network, and interacts with the synaptic tags to modify weights. That is:

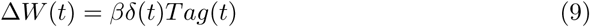

where *β* is the learning rate. Note that the two forces that determine weight updates are the RPE and the synaptic tags. The RPE signal assures that once the model accurately predicts rewards, the resulting *δ*(*t*) = 0 and the weights remain unchanged, which allows the model to converge on an on-policy solution. The synaptic tags, on the other hand, solve the credit assignment problem by means of attention gated feedback: units in the hidden layer whose activity had a larger influence on the Q-value of chosen actions receive stronger feedback and form stronger tags, whereas units that did not contribute to the selected action will not have weight updates. Previous work has established the relation between the AuGMEnT learning rule and error-backpropagation [36].

In all simulations, the model was trained using the same, general principles that are in line with typical animal learning. Changes in the environment, and the reward that was delivered, depended on the external actions of the agent, whereas internal actions that pertain to WM updates were never directly rewarded. Trials were aborted without reward delivery whenever the model selected an incorrect motor response. Reward could be obtained twice in a trial. First, all tasks required the agent to perform a ‘default action’ throughout the trial (such as maintaining gaze at a central fixation point or holding a response lever) until a memory-informed decision had to be made. We encouraged the initial selection of this action by offering a small ‘shaping’ reward (*r* = 0.2) for selecting this action at the first time step. At the end of a trial, if the correct decision was made in response to a critical stimulus, a large reward (*r* = 1.5) was delivered. In our model assessments, trials were only considered ‘correct’ when both rewards were obtained.

Although not all inputs and computations were strictly necessary or useful in every task, the network architecture, parameter values and the representation of inputs were kept constant across simulations; Across tasks, we modified only the external action module to represent the valid motor responses for the different tasks.

## Results

### Task 1: Delayed Match to Sample

Arguably one of the most straightforward WM tasks is the Delayed Match-to-Sample (DMS) task, where an agent is presented with one stimulus, and has to determine whether a second stimulus, presented after a delay without a stimulus, is the same or not. Here, we show that random, untrained encoding projections can be used to solve this task and that the solution generalizes to stimuli that the agent has not observed before. We trained the agent on a simple DMS task, where it was sequentially presented with a fixation cross, a probe stimulus, another fixation cross, and a test stimulus that would either match the probe or not (Fig. 2A,B). Stimuli consisted of unique binary patterns of six values (See Fig. 2B for two example stimuli). One additional seventh input was used to signal the presence of the fixation dot. The agent had to withhold a response until the test stimulus appeared and it then had to make one of two choices to indicate whether the test stimulus matched the probe (we used a ‘leftwards/rightwards’ saccade for ‘match/mismatch’). We modeled a total of 750 networks with randomly initialized weights. During initial training, the probe-and test-stimuli were chosen from a set of three unique stimuli (Set 1). Once performance had converged (¿ 85% correct trials), the stimulus set was replaced by a set of three novel stimuli (Set 2) This process was repeated until performance had converged for six sets of stimuli.

In these and all other simulations, we will report convergence rates based on all trials including those with exploratory actions.

Fig. 2 illustrates how the network solves an illustrative trial where probe and test do not match. The three types of inputs provided to the network during the trial are depicted in Fig. 2C, with time input, sensory input, and match input depicted from left to right. Activity of each time cell unit peaks around a unique time point, together conveying a drifting representation of time. The sensory input cells represent the stimuli presented to the agent. We have depicted three curves to summarize the activity corresponding to the fixation dot, probe and test stimulus, but we note that in mismatch trials, the two representations still usually have partially overlapping input units. The activity in the match nodes conveys the result of comparing the content of each memory block to currently presented stimulus: “Match 1” and “Match 2” for the comparison with the content in ‘memory block 1’ and ‘memory block 2’ respectively. The example agent learned to store the probe stimulus in ‘block 2’ and to correctly maintain the probe item throughout the trial, so that match signal from this block could be used for the final match-versus mismatch decision. The example trial is a mismatch, but the right panel of Fig. 2C also illustrates the activity of the match node on a trial in which the test matched this probe (dashed line and cross).

Output layer activity on this example trial is depicted in Fig. 2D. Activity in this layer approximates the value associated with the different gating-and motor actions, which influence the RPE and thereby drive learning. The total estimated Q-value, i.e. the sum Q-values of selected actions in the two modules is plotted in the left panel. They reasonably approximate the real Q-values, which sufficed for an adequate policy – they would become even more accurate after further training. The other two panels of Fig. 2D show the Q-values for all possible actions at each time step, separately per module. Note that the individual Q-values in these modules do not allow for straightforward interpretation, as they are free to vary as long as their sum provides a good Q-value estimate. In practice, however, we found that the two modules evenly contributed to the Q-value estimate.

To examine whether the policy acquired by the agents generalized to novel stimuli, we assessed the number of trials that an agent required to converge after each switch to a new set. The results in Fig. 3 illustrate that agents were able to generalize across stimulus sets. Convergence on the first set was relatively slow (Fig. 3A): the median number of trials needed for convergence was ≈ 12, 700 trials, with 95% of the agents converging within 4,379 – 46,724 trials. Already after the first switch, convergence occurred much faster, after a median of 1,066 trials (95% within 212 – 5,288, trials). With each subsequent switch, the agents displayed further generalization, and the median trials needed for convergence on set 6 was only 343.5 trials (95% within 90 – 1,994 trials. We note that 85 trials is the absolute minimum number of trials before any agent could reach our criterion of 85% accuracy).

**Fig 3.**
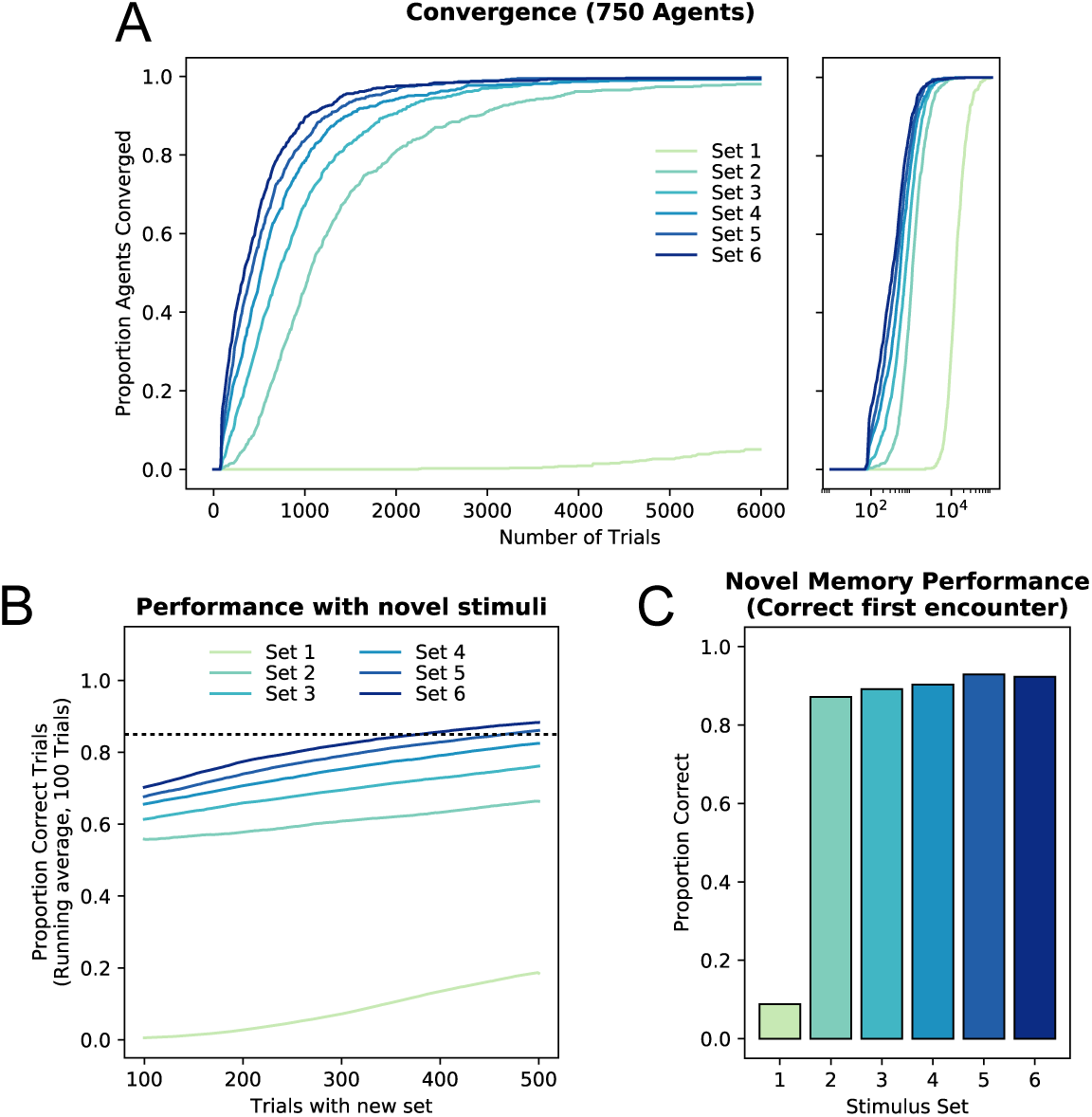
Performance and training on the Delayed Match to Sample task with novel stimuli. **A:** Convergence rates across 750 agents (left: performance on first 6000 trials with each new stimulus set; right: convergence on all sets, with log time scale). On the first set, convergence is relatively slow, but on subsequent sets, agents learn much faster. The convergence rates keep increasing with each new set. **B:** Performance with new stimuli immediately after a switch increases with each switch, indicating that the agents generalize the task to new stimulus sets. **C:** Accuracy for the first encounter with a novel test stimulus, i.e. on the first trial in which the model maintained fixation until the test stimulus was presented. Note that accuracy is 87.1% on Set 2, after the first stimulus switch. The agents then further generalize the rule across contexts, because accuracy is 90% or higher for all subsequent set switches.

We next assessed performance in the first 100 to 500 trials with each new set, to explore how fast the agents learned the task with novel stimuli (Fig. 3). Initial performance on Set 1 (after 100 trials) was near chance level, which was approximately 1% correct in this task (four consecutively correctly selected actions chosen from three response options). Performance gradually increased, reaching 18.5% accuracy within the first 500 trials. Following the first switch (to Set 2), performance did not drop back to chance: rather, agents immediately performed 55.8% correct on the first 100 trials, and were 66.3% correct after 500 trials. On each subsequent set switch, immediate performance with never-before seen stimuli kept increasing, with performance at 70.3% for the final set. On the final two sets, criterion performance (85%) was acquired within 500 trials. These results suggest that agents were indeed able to generalize the acquired policy to novel contexts, although each set switch still required some additional learning.

We suspected that one important reason why the model failed to immediately generalize on new sets, might have been that agents broke fixation for novel stimuli. Note that a completely novel input pattern makes use of connections that have not been used before in the task, which could due to their random initialization trigger erroneous saccades. To account for such errors, we also assessed the accuracy of agents on the first trial in which they encountered a novel probe and maintained fixation until the test stimulus. We observed an average accuracy of 87.1% across agents on their first encounter with a novel stimulus from Set 2. This accuracy score also increased for subsequent sets, with an average accuracy of ≈ 92.6% correct for the first encounters with stimuli from Set 5 and 6. Thus, the model learned the matching task in a manner that allows almost immediate generalization to new stimulus sets: the vast majority errors in later sets were caused by fixation breaks.

### Task 2: 12-AX

We next examined the performance of WorkMATe on the 12-AX task, a task that was used to illustrate the ability of PBWM to flexibly update WM content. The 12-AX task is a hierarchical task with two contexts: ‘1’ and ‘2’. In the task, letters and numbers are sequentially presented, and each require a ‘go’ or a ‘no-go’ response. Whenever a ‘1’ has been presented as the last digit, the ‘1’-context applies. In this context, an ‘A’ followed by an ‘X’ should elicit a go response to the ‘X’, whereas every other stimulus requires a no-go response. When a ‘2’ is presented, the second context applies: now only a ‘B’ immediately followed by a ‘Y’ should elicit a go response. Agents must separately maintain and update both the context (‘1’ or ‘2’) and the most recently presented stimulus, in order to make the correct go response to the imperative stimuli ‘X’ or ‘Y’.

Human participants can do this hierarchical task after verbal instruction, but to acquire the rules that determine the correct response solely through trial and error learning poses a challenge. PBWM learned this task using a complex combination of reinforcement learning, supervised learning, and unsupervised learning techniques [56, 58, 80], but [55] showed that agents can also learn this task using a simpler *SARSA*(*λ*) reinforcement learning scheme. To our knowledge, no data have been published on humans or other primates learning a task of this complexity through reinforcement learning alone.

Here, we used a trial-based version of the task, where on every trial a sequence of symbols with unpredictable length is presented, which ends with an ‘X’ or a ‘Y’. During this sequence, the agent had to respond as outlined above. Given the complexity of the task, we trained the agents through ‘curriculum learning’ [50, 81, 82], a training scheme in which trials were organized into ‘levels’, which gradually increased in difficulty. Once an agent showed sufficient performance on a level, training for the next levels commenced. Example sequences at different difficulty levels are shown in Fig. 4A. Key to curriculum learning is that trial types from previous, easier levels are also presented in order to prevent unlearning of the simpler cases. In our curriculum, 50% of the trials were always of the highest difficulty level, and the other 50% simpler cases drawn from one of the previous levels with equal probability for all previous levels. The difficulty was increased when performance on the last 100 trials was over 85% correct.

**Fig 4.**
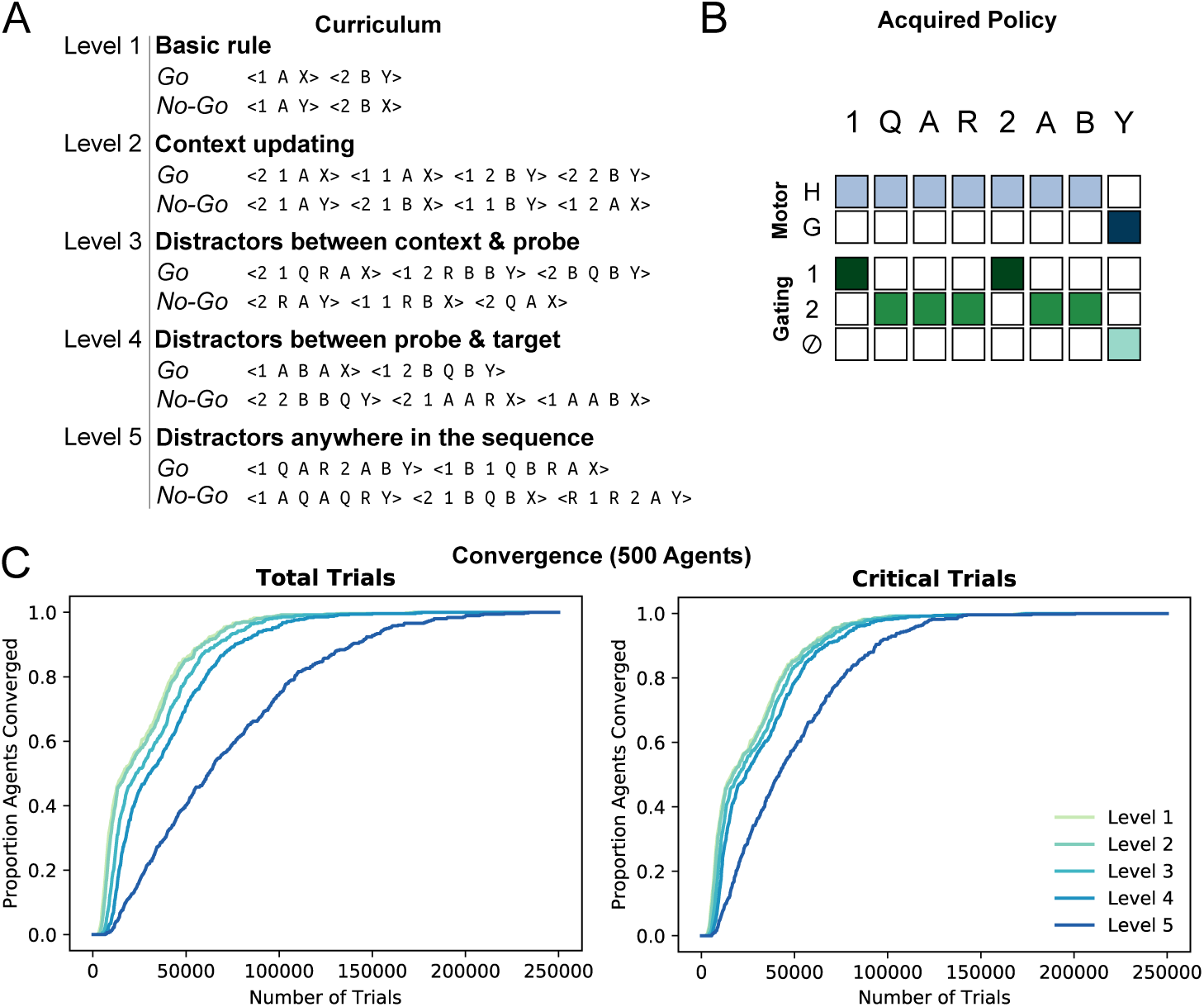
Training on trial-based 12AX. **A:** The curriculum used to train the agent, with example trial sequences to illustrate the difficulty levels. As soon as the agent performs correct on 85% of the trials, a higher difficulty level is introduced and presented on 50% of the trials (critical trials), with the other 50% sampled from the lower levels. **B:** Policy on an example trial (cf. Fig. 2E), acquired by an illustrative model agent converging on the highest difficulty level. The agent correctly updates memory content on each stimulus, but is only rewarded on the basis of its final motor action in response to the target symbol (X/Y). This agent stored the task context (1/2) in the memory block 1, and stored the last seen stimulus, target or distractor, in block 2. **C:** Cumulative Histogram from 500 agents depicting the number of trials needed for convergence on each difficulty level. Training on higher difficulty levels does not start until lower levels have been learned. The graph on the right depicts convergence rates considering only the trials drawn from the highest difficulty.

This trial-based curriculum not only facilitated training, but it also had another benefit over previous approaches to train 12-AX [1, 55, 83]. In previous implementations, the imperative X/Y stimulus always occurred at one of a few critical moments after the context rule, whereas here we intermixed sequences of very different lengths. Without this variation, we found that models could meet the convergence criterion on the basis of timing alone, without fully acquiring the task rules. In the current curriculum learning scheme, the agents truly solved the task, applying the appropriate storage policies to all difficulty levels and trial lengths.

All 500 agents converged and were able to accurately perform the task at the highest difficulty level. The policy acquired by one of these agents is depicted in Fig. 4B, which illustrates an example trial at the highest difficulty. Throughout the sequence, the agent selected the ‘hold’ action, while it updated each last-presented stimulus, encoding these into block 2. However, stimuli denoting the rule context (1/2) were encoded into memory slot 1, and only updated when the context changed. Once presented with the imperative stimulus (‘Y’, in this case), this gating policy allowed the agent to use the memory of the current context (‘2’) and the previous stimuli (’B’) to decide to yield a correct ‘go’ response.

Convergence rates for this task are depicted in Fig. 4C. Despite the complexity of this task, all agents reached criterion performance, within a median of ≈62,000 trials (95% range 11,566, – 180,988). A large proportion of these trials were repetitions of easier levels, and the number of critical (final level) trials before convergence was lower, with a median of ≈42,000 trials (95% 8,700 – 121,208). Thus, the model was able to acquire the rules of complex, hierarchical task which requires flexible gating of items into and out of WM, based only on relatively sparse rewards that were given only at the end of correctly performed trials.

### Task 3: ABAB ordered recognition

In a series of elegant studies, Miller and colleagues [2, 3, 15, 84], reported data from macaques trained in tasks in which multiple visual stimuli needed to be maintained in WM. For example, in the ‘ordered recognition task’, the monkey was trained to remember two sequentially presented visual stimuli (A and B), and to report whether the stimuli were later presented again, in the same order. On match trials the same objects were repeated (ABAB), and the monkey responded after a match to both objects, i.e on the fourth stimulus in the sequence. There were mismatch trials in which the first or the second stimulus was replaced by a third stimulus C (ABAC or ABCB) as well as mismatch trials with the same stimuli (A and B), but in reverse order (ABBA). In case of a mismatch, the monkey waited until the A and B were shown in the correct order as the fifth and sixth stimuli (e.g. ABACAB), and thus responded to the sixth stimulus. In each recording session, three novel visual stimuli were used to form the sequences, where each of these stimuli could take on the ‘role’ of A, B or C on any trial.

This ordered recognition task requires selective updating and read-out of memories in a way that shares features with the 12-AX and DMS tasks from the previous sections. As in the 12-AX task, two stimuli need to be maintained and updated separately, and the task goes beyond simply memorizing two items: the order of stimuli also needs to be stored and determines the correct action sequence. As with the DMS task, monkeys reached reasonable accuracies, even though novel stimuli were presented in each session, implying that they could generalize their policy to new stimulus sets.

We tested WorkMATe on this ordered recognition task. We trained 750 model agents, randomly selecting stimuli from the same set as we had used for the DMS simulation described above. Half of the trials were ‘match’ sequences, and the other half consisted of the three possible mismatch sequences, in equal proportion. Criterion performance was defined as an accuracy of at least 85% on the last 100 trials, with an added requirement of at least 75% accuracy on the last 100 trials in each of the four conditions. In the ‘static’ training regime, we kept the three selected stimuli identical for an agent throughout a training run. In the ‘dynamic’ regime, the three stimuli were replaced by three new randomly selected stimuli after 3,000 trials. This meant that each of the three stimuli took on the role of A, B or C approximately 1,000 times before they were replaced by a new set.

The convergence rates for the static regime are plotted as solid lines in Fig. 5A. The agents learned the full task after a median number of ≈106,000 trials (95% of the agents between 25,880 – 856,868 trials). Under the static regime, we found that learning the overall task was primarily hindered by the condition ‘Mismatch 1’ (ABCB). Convergence on this condition typically took much longer (median: 86,390) than on the other conditions (medians: 3,128, 24,076, 28,858 trials for ‘Match’, ‘Swap’, and ‘Mismatch 2’ respectively). The increase in complexity under the dynamic regime caused a total training time that was five to six times longer (Fig. 5A, dashed lines) than in the static regime, with convergence after a median of *≈*641,000 Trials (95% of the models converged within 139,907 – 3,797,200 trials). Interestingly, compared to the static regime, initial convergence was comparatively quick on each of the mismatch conditions, within a median of ≈13,000 trials (75% correct). The reason for this is that many agents initially learned to withhold their response until the end of the trial, but did not learn to store or update the appropriate stimuli in WM. Even though all mismatch conditions initially converged rather quickly, we noticed that during training, increases in ‘Match’ condition performance were often paired with decreases in performance on the ‘Mismatch 1’ condition.

**Fig 5.**
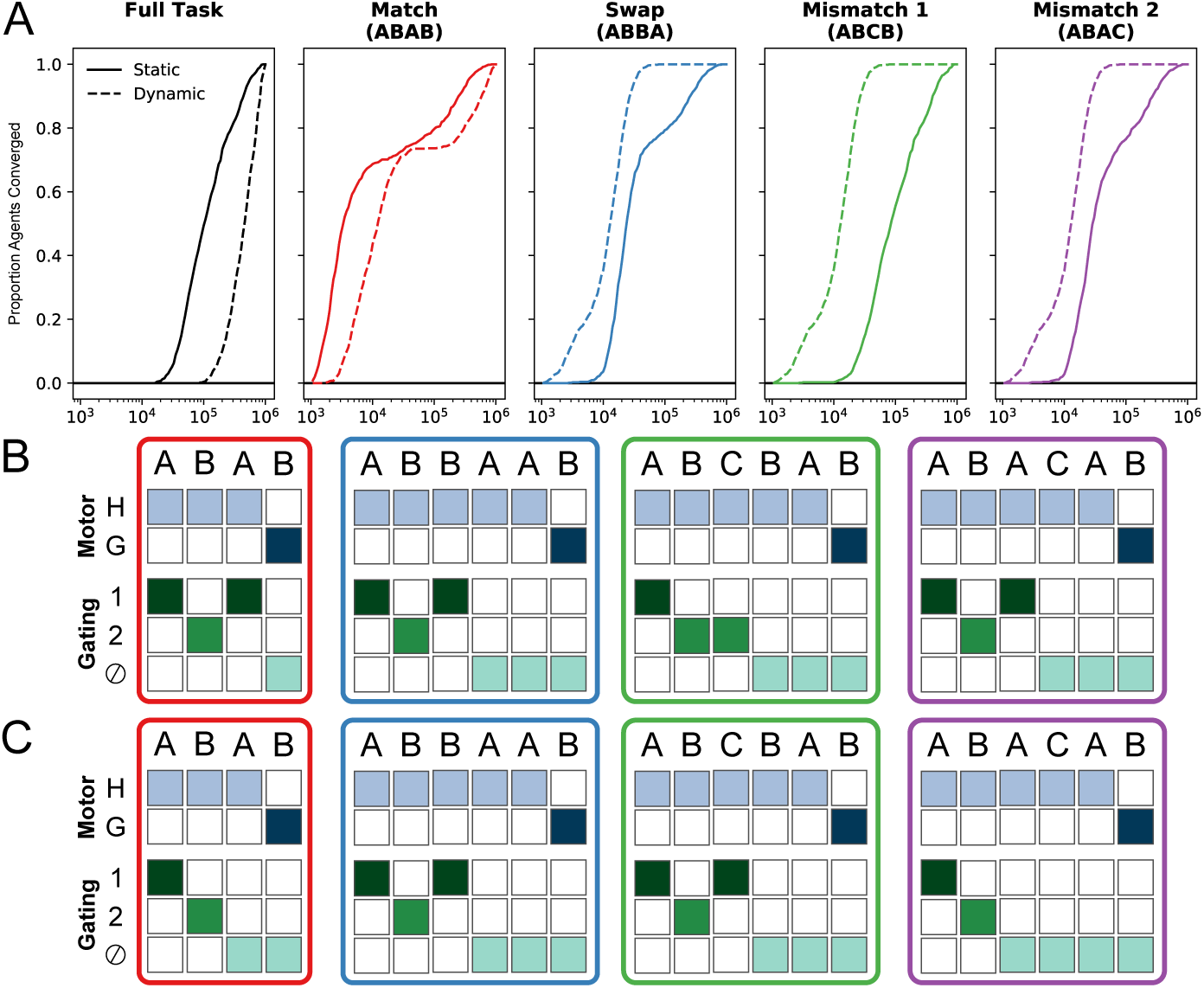
The ABAB ordered recognition task. **A:** Convergence of 500 agents on the full task (black traces) and on different conditions separately (colored traces) under two training regimes: static (same stimuli used throughout training) or dynamic (new stimulus sets after each 3,000 trials). In both regimes the task is typically learned in approximately 10^5^ trials, but convergence varies across conditions. Note the logarithmic time axis. **B:** The ‘memorized mismatch’. **C:** The ‘memorized storage time’ policy. Both policies reflect generic, common solutions found amongst converged agents, and are discussed in the main text. Both are plotted following Fig. 2E.

We qualitatively investigated the policies of converged agents to explore why the ‘Mismatch 1’ posed such a challenge for the model. Note that on trials from the other conditions (‘Match’, ‘Swap’, and ‘Mismatch 2’, which together make up 83.3% of all trials) the correct response can be determined based on relatively simple inferences: The agent merely has to learn to encode the second stimulus (B), and maintain it for two time steps, and utilize its time cell input to identify the fourth and sixth stimulus presentation. Then, if the stimulus at *t* = 4 matches the stimulus that was encoded at *t* = 2 a Go-response is needed, otherwise it is to be held until *t* = 6. The ‘Mismatch 1’ condition, however, demands complex memory management. The agent must store both the initially presented A and B, detect the mismatch at *t* = 3, and somehow convey this mismatch in a manner that prevents responses to the matching stimulus (B) at *t* = 4. However, in the present architecture, WorkMATe has no way to encode this mismatch, so the agent is not capable of such meta-cognition.

Nevertheless, agents typically found a solution that fell into one of two classes: In both solutions, the first two stimuli (A/B) were separately encoded in the two memory blocks. The first solution, which we call the ‘memorized mismatch’ strategy (Fig. 5B), essentially followed the following rule: If the stimulus at *t* = 3 does not match either stimulus in memory – and the trial must therefore be of the ‘Mismatch 1’ condition – the agent replaced the ‘B’-stimulus in memory with the ‘new’ stimulus C. As a result, stimulus ‘B’ at *t* = 4 no longer matched any stimulus in memory, which led the agent to withhold a response. A second solution, the ‘memorized storage time’ strategy, made use of the temporal information encorporated in the memory representation of the stimulus. In this strategy, the key step was that if the stimulus at *t* = 3 did not match the ‘A’, the initial A-stimulus was overwritten in memory. At *t* = 4, the correct decision could then be made by only responding if stimulus matched the ‘B’ in memory, and if the other memory block contained temporal information from the first time step.

To conclude, these simulations demonstrate that WorkMATe can acquire complex control over WM content, in order to appropriately solve complex hierarchical tasks with dynamically switching stimulus contexts – again, solely on the basis of reinforcement signals.

### Task 4: Pro-/Anti-saccade Task

To compare WorkMATe to its ‘gateless’ predecessor AuGMEnT [36], we simulated agents learning the delayed pro-/ antisaccade task, a classic task in both human and non-human primate memory research, and on which AuGMEnT was also trained and evaluated. The task (Fig. 6A) requires an agent to maintain fixation at a central fixation point. The agent should encode the location of a peripheral probe and memorize it during a delay. Trials with a black fixation point are pro-saccade trials and when the fixation point disappears, the agent makes a saccadic eye movement to the remembered location of the probe. On anti-saccade trials, the fixation point is white and now the agent has to make an eye movement in a direction opposite to the remembered cue location, after the memory delay.

**Fig 6.**
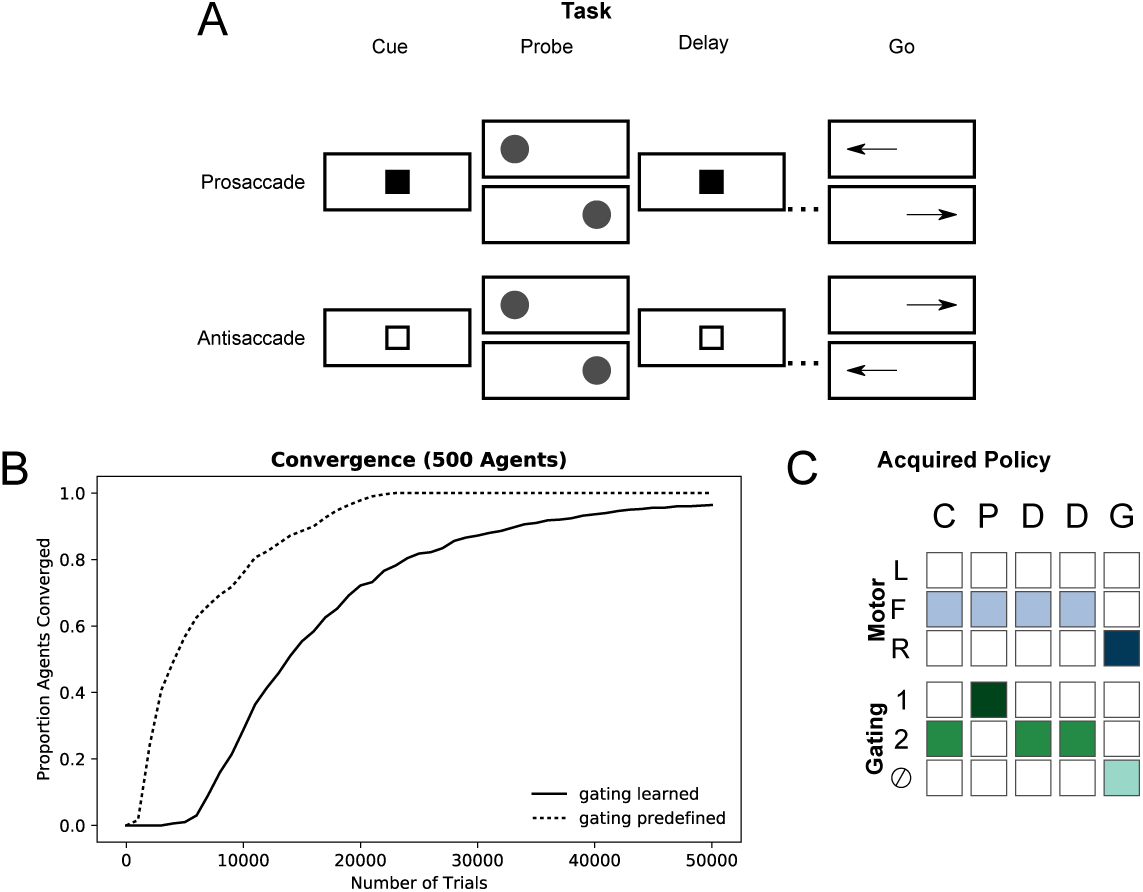
A: Illustration of the four conditions in the prosaccade/antisaccade task. The agent has to memorize the location of the probe, and make a pro-or anti-saccade after a delay, dependent on the trial type indicated by the cue (white or black fixation point). The agent thus has to integrate the information throughout the trial, and make an “exclusive or” decision upon presentation of the go-signal. Of note, the gating policy in this trial, depicted in **C**, is applicable in each of the four conditions in this task. **B:** Convergence rates for 2 × 500 simulated agents of two different types. The solid line depicts convergence with WorkMATe. The dotted line depicts performance with a modified version of the model, where the gating policy is not learned, but correctly predefined and fixed beforehand. **C:** Policy (cf. Fig. 2E) of an example agent after convergence, during an antisaccade trial with a ‘left’ probe. This gating policy applies to all trial conditions.

We trained 500 instances of our network and all learned the task (¿ 85% correct) within 100,000 trials (Fig. 6B, solid line). The median number of trials was ≈ 15,000 (95% 6,835 – 56,155 trials). This convergence rate is faster than that of monkeys, who typically learn such a task only after several months of daily training with ≈ 1,000 trials per session. However, training took approximately three to four times *longer* than with the original AuGMEnT architecture. There are several differences between AuGMEnT and WorkMATe that could account for this. For example, the parameters governing Q-learning were not optimized for WorkMATe, but adopted from AuGMEnT to facilitate comparison. The most critical difference between models, however, is that the gated memory store which is the core of the WorkMATe model, was overly flexible for this task. The gateless AuGMEnT architecture encoded all relevant stimuli into its memory so that an accumulation of relevant information was available at the ‘go’ signal. The WorkMATe architecture first had to acquire an appropriate gating policy (Fig. 6C), to make sure that the correct decision can be made based the fixation color and probe location on the ‘go’ display when no information is available anymore. Notably, the gating policy can be the same for all conditions: if cue and probe are separately available in memory, a correct decision can be made.

To examine if the added complexity of learning a gating policy could account for the difference in learning speeds between WorkMATe and AuGMEnT, we trained a new set of ‘gateless’ agents on this task. These agents were identical to WorkMATe, except that the gating actions were, from the start, predefined to match those depicted in Fig. 6C. With this setup, the complexity was comparable to that of the AuGMEnT architecture. Indeed, convergence rates for these gateless agents (median number of trials ≈ 5,000; 95% 2,076 – 20,334 trials) were very similar to those for AuGMEnT, and were approximately three times faster than those with gated WorkMATe (Fig. 6B).

These simulations highlight the strengths and weaknesses of ‘gateless’ and ‘gated’ memory architectures. Simpler, gateless models that project all stimuli to memory suffice for tasks like pro-/antisaccade task. These tasks do not require selective updating of memory representations, nor do they contain distractor stimuli that interfere with the memory representation. On the other hand, gating is essential for tasks in which access to WM needs to be controlled in a ‘rule-based’ fashion. In both the ABAB ordered recognition task and the 12-AX task, a stimulus’ access to memory is contingent on other items that are presented in the history of the trial. We envisage that both types of WM, gated and ungated, might exist in the brain, so that the advantages of both strategies can be exploited when useful.

## Model Stability

Our simulations demonstrate that the WorkMATe model is able to learn accurate performance across a range of popular WM tasks. Across these simulations, we have kept the model architecture and parameters constant, including the learning parameters *β*, which scales the synaptic weight updates, and the SARSA learning parameter *λ*, which determines the decay of synaptic tags, through the relation *α* = 1 − *λγ*. Our choice of values for these parameters was mainly motivated by consistency with the original AuGMEnT model. In order to explore to what extent WorkMATe’s performance remains stable across variability in these parameters, we ran a grid-search exploration of the parameter space for different values of *λ* and *β*.

For this grid search we used versions of the tasks defined for the simulations above. For the DMS task, we used only three stimulus sets. For ABAB ordered recognition task we only ran the ‘Static’ learning regime (solid lines in Fig. 5A). For 12-AX, we again used curriculum learning, and count only the ‘critical’ trials at the highest difficulty level (cf. Fig. 4C). The pro-/antisaccade task was ran as-is (solid line in Fig. 6B). We assessed all combinations of *β* = [0.05, 0.10, 0.15…1.0] and *λ* = [0.1, 0.2, 0.3…0.9], and for each parameter combination, we ran 100 model instances. The simulations were ran on the Peregrine High Performance cluster of the University of Groningen. Per task, each model instance was allotted the same amount of wall clock time. Assuming comparable performance across all cores this implies a similar maximum number of iterations (trials) held for these tasks. The maximum number of iterations in each task was ≈ 1, 870, 000 in the Delayed Match to Sample task, ≈ 1, 700, 000 critical trials in the 12-AX task, ≈ 790, 000 trials for ABAB ordered recognition, and ≈ 500, 000 in the Pro-/ Antisaccade task. The number of iterations reported in Fig. 7 are the median number of iterations computed across all runs in which convergence was reached. In general, we found that model runs with a high *β* had relatively low convergence rates, an effect that was particularly pronounced for the ABAB task. To yield better insight into model stability for this task, we ran additional simulations where we varied *β* at a more fine-grained scale *β* = [0.025, 0.05, 0.075…1.0].

**Fig 7.**
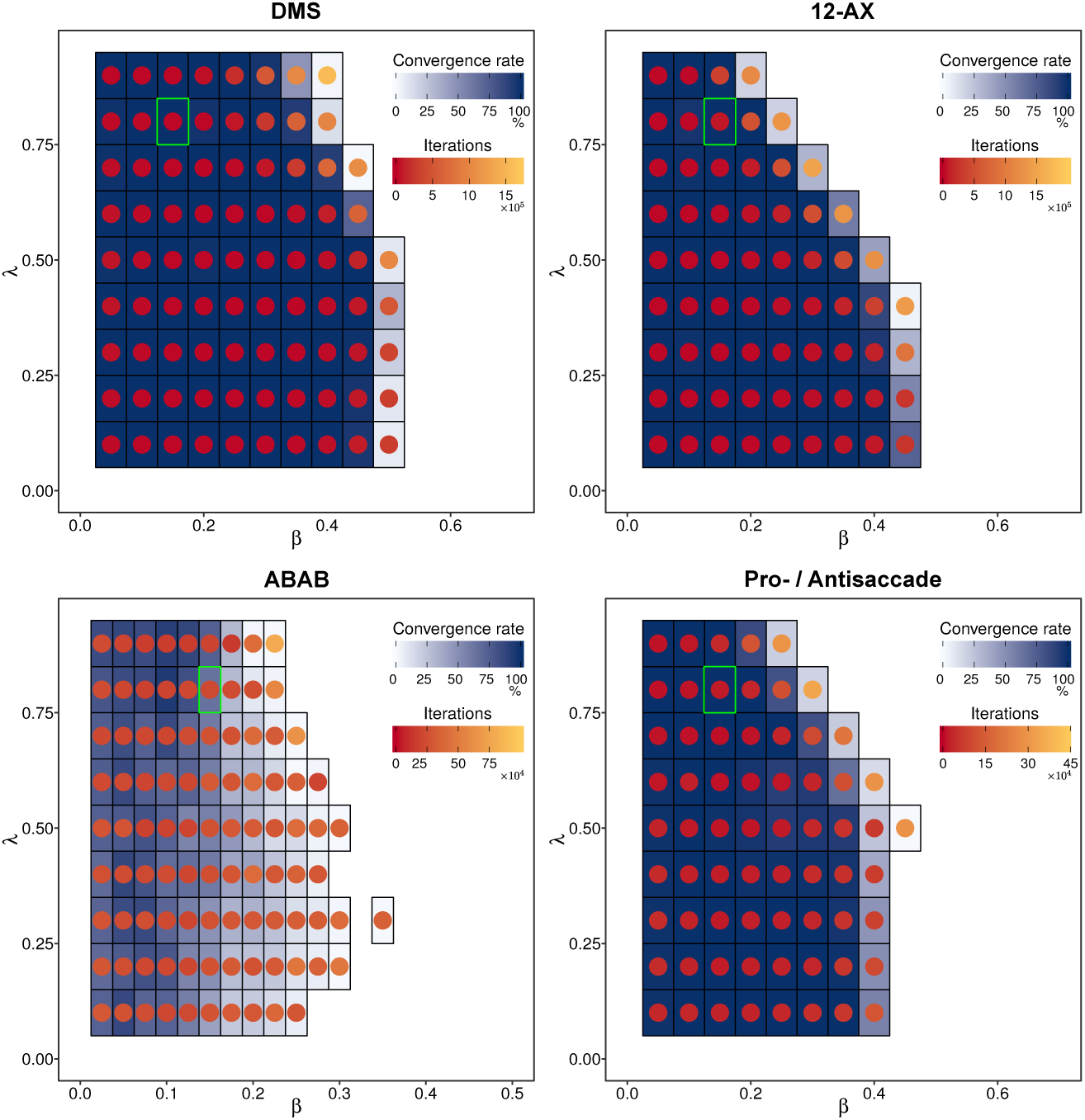
Model stability across the four different tasks. Each tile represents a parameter combination. The blue shading of the tiles indicates the convergence rate, and the color of the dots the median number of iterations for convergence. Note that the color axis for iterations is different for each task, and the x-axis is different for the ABAB ordered recognition task (bottom left). The green outline indicates the single parameter combination that was used in all simulations in the preceding sections. It can be seen that model performance is largely independent of values for *λ*, and that lower *β* values were generally associated with faster convergence.

The results are depicted in Fig. 7. Across all tasks, a similar pattern was found: performance was rather robust across a range of values for *λ*, and more sensitive the precise value of *β*. Parameter *λ* determines the decay rate of the synaptic tags in the neural network that are formed once an action is selected. Notably, variations in this parameter impacted performance most the Pro-/ Antisaccade task and 12-AX. These were tasks where reward delivery could be furthest removed from the actions that led to a possible reward. For example, during 12-AX, committing the context rule ‘1’ to memory is necessary, but might only lead to reward much later.

With regards to *β*, the results suggest that too high learning rates are detrimental for WorkMATe. Too high values for the learning rate are generally harmful for convergence in neural networks, and for WorkMATe this might have been extra detrimental due to the all-or-none gating policy in the model. Gradual weight changes could therefore lead to sudden changes in the gating policy, altering the model’s state space. Large learning rates can therefore prevent convergence by rendering previously learned state-action pairings irrelevant. Although these sudden changes also occur with lower *β* values, they are less frequent so that the models can adapt. Of note, the influence of *β* and *λ* on learning was similar to that observed with previous models [36, 55]. We conclude that there are large regions of the parameter space with successful and consistent performance in all four tasks. Within these regions the performance of WorkMATe is robust and stable.

## Discussion

We have presented WorkMATe, a neural network model that learns to flexible control its WM content in a biologically plausible fashion by means of reinforcement. The model solves relatively basic WM tasks like delayed match to sample and delayed Pro-/ Antisaccade tasks, but also more complex tasks such as the hierarchical 12-AX task and the ABAB ordered recognition task. Furthermore, we show that the agent can learn gating policies that are largely independent of the stimulus content, and applies these policies successfully to solve tasks with stimuli that were not encountered before. Thus, WorkMATe exhibits a number of crucial properties of WM: trainability, flexibility, and generalizability.

The terms ‘working memory’ and ‘short-term memory’ have often been used interchangeably in the cognitive sciences, even though the term ‘working memory’ was popularized to place additional emphasis on the capability of the brain to flexibly regulate and update memory content given task demands [85]. Many previous models of WM (e.g. [7, 9, 13]) focus on storage of items and their retrieval. In the present work, the focus was on learning to use and update memory content according to possibly complex task requirements. WorkMATe models acquire gating-and updating policies that implement a type of ‘symbolic’ memory control: In many of our simulations, the acquired gating policy can be interpretated as a set of production rules that are applicable to all conditions of the task, independent from the precise identity of the stimuli. Previous studies have noted that the gap between traditional artificial neural network architectures and symbolic systems is one of the great challenges to be overcome by artificial intelligence [86]. Previous neural network models that attempt to implement a similar approach to memory control have relied on predetermined, hand-coded sequences of memory operations hard-coded into the model ([87–89], but see [50]). Here we show, for the first time, that such control over WM can be acquired in a neural system by means of biologically plausible reinforcement learning.

These strengths of WorkMATe originate from the combination of design features of previous neural network models of WM. We sought to overcome problems faced by action-oriented models such as PBWM, LSTM and AuGMEnT by combining the AuGMEnT learning rule with a memory circuit inspired by more generic memory models. WorkMATe can store arbitrary representations, and has the built-in capacity to compute the degree of match between the representations in memory and incoming sensory information. The generality of the model follows from our finding that it is unnecessary to first learn specific memory representations, and that instead a fixed, random projection for encoding suffices. The properties of such an encoding scheme have been explored before [76, 77], indicating that this is a functionally rich approach that can be applied to a range of memory tasks. Our simulations with the pro-/antisaccade task demonstrate that such random feedforward encoding suffices for at least some tasks where the relevant features are given as feedforward inputs to the model. It seems likely however, that it will be insufficient for other tasks, in which the memoranda require specific and non-linear combinations of inputs. Recently, [78] proposed a working memory storage architecture that was defined by two separate layers of neurons: a structured, sensory layer with separate pools for separate items, which projected to a shared ‘unstructured’ layer via random recurrent connections, with their only constraint being that excitation and inhibition were balanced for each neuron. The resulting architecture could also store arbitrary representations, and gave rise to capacity limits and forgetting dynamics that are also observed in humans. Future work might explore how WorkMATe might also benefit from a more sophisticated memory maintenance architecture, be it a multi-layer subsystem or one with recurrent connections to the sensory inputs, while still allowing for the generic, built-in matching computations. Indeed, it is this matching process that endows WorkMATe with the flexibility of dealing with stimuli that were not previously encountered by the model.

WorkMATe makes several simplifying assumptions that touch on contended topics in WM research, and therefore require further discussion. First, all our simulations made use of two, independently maintained memory blocks to store content, which proved sufficient for these tasks. There is an ongoing debate regarding the storage capacity limits of WM, and to what extent these speak to the functional organization of items in memory. Two opposing views are slot-based models [69], which state that storage is limited by a discrete number of ‘slots’ in memory, and resource-based models which propose that there is no limit on the number of items that can be stored, but the total fidelity is limited by a certain amount of ‘resources’ [71, 72, 90]. Our memory circuit most closely aligns with the slot-based view, but it is conceivable that similar results might have been obtained with a resource-based implementation. However, an approach with independent memory blocks allows for independent matching, gating and updating of memoranda, for which resource-based architectures would require additional assumptions (See [68] for one potential approach). Furthermore, while two blocks sufficed for the tasks simulated here, it is conceivable that other tasks performed by humans might require more blocks, because the capacity limit of WM in humans is larger than two items [70, 91, 92]. Our focus, however, was not on slot-based architectures or the exact limitations, but on learning to flexibly control multiple representations.

A second simplifying assumption that we have made here is that matches between sensory-and memory representations are computed automatically and in parallel. Whether multiple memory objects can be ‘matched’ at once by a single percept is a topic of debate in cognitive psychology [93–97]. The tasks that we have chosen to focus on here take place at relatively slow speeds, which would allow for serial comparisons, but previous research has shown that at high speeds, matching multiple memory targets comes at a cost [98]. A serial comparison circuit might introduce additional control operations to determine which representation should be prioritized for matching. Here, we refrained from simulating such additional operations. Related to this, some models such as LSTM can also gate WM *output*, in addition to the input. These might come into play in task-switching setups, where multiple goals need to be maintained but only one should drive behavior [99–104], and in sequential visual search tasks where multiple items may be held in WM but only one drives attentional selection [95, 105–110]. Recordings in Macaque PFC suggest that sequential search tasks, which require such ‘prioritization’, of one memory item over another are characterized by elevated cortical representation of the prioritized stimulus in preparation of search [2, 3, 15]. Future extensions of WorkMATe might investigate tasks that could benefit from such output gating operations, and whether they can be learned through plasticity rules related to those studied here.

Interestingly, not every task benefited from a gated memory. Notably, training on the pro-/antisaccade task actually took 3-4 times longer with the gated model than with a model without these gates. This is important, as it shows that for certain tasks, it may indeed be beneficial to merely accumulate relevant information into memory and learn a policy that relies on these accumulated representations. These types of memory tasks are actually more akin to perceptual decision-making tasks, which require an agent to aggregate information until a threshold is reached that triggers a decision [111, 112], rather than to flexibly store, update and maintain memory representations. This qualitative dissociation between different types of tasks might warrant a model that is comprised of separate routes to a decision: one relying on the automatic integration of relevant information, and one describing a more controlled process that stores and updates information as variables to be used in a task [113].

Intriguingly, recent work by [114] used a very different approach to WM modeling but arrived at a similar conclusion. They trained networks with recurrent pools of units using a supervised stochastic gradient descent algorithm, in order to solve different working memory tasks. Their findings indicated that functionally very different types of circuits were used to solve different working memory tasks, dependent on the degree of dynamic control and updating that the task demanded. Our simulations similarly suggest a dissociation between tasks that rely on the simple integration of relevant information whereas others may benefit from, or even demand additional levels of control. We may therefore use models like WorkMATe to predict more precisely which tasks will rely on flexible, controlled memory, and which tasks can be solved without the necessity for flexible control structures (the actual ‘work’).

## Conclusion

To conclude, we have presented a neural network model of primate WM that is able to learn the correct set of internal and external actions based on a biologically plausible neuronal plasticity rule. The network can be trained to execute complex hierarchical memory tasks, and generalize these policies across stimulus sets that were never seen before. We believe this to be an important step towards unraveling the enigmatic processes that make WM ‘work’: that is, be used as an active, flexible system with capabilities beyond the mere short-term storage of information.

## Supporting information

**S1 Table.** Model paramaters. Parameters used in all simulations (unless stated 756 otherwise).

| Symbol | Name | Value |
| --- | --- | --- |
| $\beta$ | Learning rate | 0.15 |
| $\gamma$ | Temporal discounting factor | 0.9 |
| $\lambda$ | Eligibility Trace decay rate | 0.8 |
| $\epsilon$ | Exploration rate | 0.025 |
| Symbol | Name | Size |
| $x$ | Total input units | 17 |
| $x_s$ | Sensory input units | 7 |
| $x_\tau$ | Time input units | 10 |
| $h$ | Hidden units | 15 |
| $S$ | Memory store (Blocks) | 2 |
| $m_i$ | Units per memory block | 14 |
| $q_{\text{int}}$ | Output q-units (internal actions) | 3 |
| $q_{\text{ext}}$ | Output q-units (external actions) | 2 or 3 (Task-dependent) |

## References

1. O’Reilly RC, Frank MJ. Making Working Memory Work: A Computational Model of Learning in the Prefrontal Cortex and Basal Ganglia. Neural Comput. 2006;18(2):283–328. doi:10.1162/089976606775093909.

2. Warden MR, Miller EK. The Representation of Multiple Objects in Prefrontal Neuronal Delay Activity. Cerebral Cortex. 2007;17(suppl 1):i41–i50. doi:10.1093/cercor/bhm070.

3. Warden MR, Miller EK. Task-Dependent Changes in Short-Term Memory in the Prefrontal Cortex. The Journal of Neuroscience. 2010;30(47):15801–15810. doi:10.1523/JNEUROSCI.1569-10.2010.

4. Naya Y, Suzuki WA. Integrating What and When Across the Primate Medial Temporal Lobe. Science. 2011;333(6043):773–776. doi:10.1126/science.1206773.

5. Brunel N, Wang XJ. Effects of Neuromodulation in a Cortical Network Model of Object Working Memory Dominated by Recurrent Inhibition. Journal of Computational Neuroscience. 2001;11(1):63–85. doi:10.1023/A:1011204814320.

6. Amari Si. Dynamics of pattern formation in lateral-inhibition type neural fields. Biological Cybernetics. 1977;27(2):77–87. doi:10.1007/BF00337259.

7. Mongillo G, Barak O, Tsodyks M. Synaptic Theory of Working Memory. Science. 2008;319(5869):1543–1546. doi:10.1126/science.1150769.

8. Barak O, Tsodyks M. Persistent Activity in Neural Networks with Dynamic Synapses. PLOS Computational Biology. 2007;3(2):e35. doi:10.1371/journal.pcbi.0030035.

9. Fiebig F, Lansner A. A Spiking Working Memory Model Based on Hebbian Short-Term Potentiation. Journal of Neuroscience. 2017;37(1):83–96. doi:10.1523/JNEUROSCI.1989-16.2017.

10. Oberauer K, Lin HY. An interference model of visual working memory. Psychological Review. 2017;124(1):21–59. doi:10.1037/rev0000044.

11. Raffone A, Wolters G. A Cortical Mechanism for Binding in Visual Working Memory. Journal of Cognitive Neuroscience. 2001;13(6):766–785. doi:10.1162/08989290152541430.

12. Jensen O, Lisman JE. Hippocampal sequence-encoding driven by a cortical multi-item working memory buffer. Trends in Neurosciences. 2005;28(2):67–72. doi:10.1016/j.tins.2004.12.001.

13. Schneegans S, Bays PM. Neural Architecture for Feature Binding in Visual Working Memory. Journal of Neuroscience. 2017;37(14):3913–3925. doi:10.1523/JNEUROSCI.3493-16.2017.

14. Downing P, Dodds C. Competition in visual working memory for control of search. Visual Cognition. 2004;11(6):689–703.

15. Siegel M, Warden MR, Miller EK. Phase-dependent neuronal coding of objects in short-term memory. Proceedings of the National Academy of Sciences. 2009;106(50):21341–21346. doi:10.1073/pnas.0908193106.

16. Miller EK, Erickson CA, Desimone R. Neural Mechanisms of Visual Working Memory in Prefrontal Cortex of the Macaque. The Journal of Neuroscience. 1996;16(16):5154–5167. doi:10.1523/JNEUROSCI.16-16-05154.1996.

17. Freedman DJ, Riesenhuber M, Poggio T, Miller EK. A Comparison of Primate Prefrontal and Inferior Temporal Cortices during Visual Categorization. Journal of Neuroscience. 2003;23(12):5235–5246. doi:10.1523/JNEUROSCI.23-12-05235.2003.

18. Rawley JB, Constantinidis C. Effects of task and coordinate frame of attention in area 7a of the primate posterior parietal cortex. Journal of Vision. 2010;10(1):12–12. doi:10.1167/10.1.12.

19. Ludueña GA, Gros C. A Self-Organized Neural Comparator. Neural Computation. 2013;25(4):1006–1028. doi:10.1162/NECOa00424.

20. Howard MW, Kahana MJ. A Distributed Representation of Temporal Context. Journal of Mathematical Psychology. 2002;46(3):269–299. doi:10.1006/jmps.2001.1388.

21. Lohnas LJ, Polyn SM, Kahana MJ. Expanding the scope of memory search: Modeling intralist and interlist effects in free recall. Psychological Review. 2015;122(2):337–363. doi:10.1037/a0039036.

22. Raaijmakers JG, Shiffrin RM. Search of associative memory. Psychological Review. 1981;88(2):93–134. doi:10.1037/0033-295X.88.2.93.

23. Howard MW, Eichenbaum H. The Hippocampus, Time, and Memory Across Scales. Journal of experimental psychology General. 2013;142(4):1211–1230. doi:10.1037/a0033621.

24. Norman KA, O’Reilly RC. Modeling hippocampal and neocortical contributions to recognition memory: a complementary-learning-systems approach. Psychological review. 2003;110(4):611.

25. Meyer T, Rust NC. Single-exposure visual memory judgments are reflected in inferotemporal cortex. eLife. 2018;7. doi:10.7554/eLife.32259.

26. Engel TA, Wang XJ. Same or Different? A Neural Circuit Mechanism of Similarity-Based Pattern Match Decision Making. Journal of Neuroscience. 2011;31(19):6982–6996. doi:10.1523/JNEUROSCI.6150-10.2011.

27. Sugase-Miyamoto Y, Liu Z, Wiener MC, Optican LM, Richmond BJ. Short-Term Memory Trace in Rapidly Adapting Synapses of Inferior Temporal Cortex. PLOS Computational Biology. 2008;4(5):e1000073. doi:10.1371/journal.pcbi.1000073.

28. Zelinsky GJ. A Theory of Eye Movements During Target Acquisition. Psychological Review. 2008;115(4):787–835.

29. Rao RPN, Zelinsky GJ, Hayhoe MM, Ballard DH. Eye movements in iconic visual search. Vision Research. 2002;42(11):1447–1463. doi:10.1016/S0042-6989(02)00040-8.

30. Hamker FH. The Reentry Hypothesis: The Putative Interaction of the Frontal Eye Field, Ventrolateral Prefrontal Cortex, and Areas V4, IT for Attention and Eye Movement. Cerebral Cortex. 2005;15(4):431–447. doi:10.1093/cercor/bhh146.

31. Lillicrap TP, Cownden D, Tweed DB, Akerman CJ. Random synaptic feedback weights support error backpropagation for deep learning. Nature Communications. 2016;7:ncomms13276. doi:10.1038/ncomms13276.

32. Richards BA, Lillicrap TP. Dendritic solutions to the credit assignment problem. Current Opinion in Neurobiology. 2019;54:28–36. doi:10.1016/j.conb.2018.08.003.

33. Whittington JCR, Bogacz R. An Approximation of the Error Backpropagation Algorithm in a Predictive Coding Network with Local Hebbian Synaptic Plasticity. Neural Computation. 2017;29(5):1229–1262. doi:10.1162/NECOa00949.

34. Scellier B, Bengio Y. Equivalence of Equilibrium Propagation and Recurrent Backpropagation. Neural Computation. 2018;31(2):312–329. doi:10.1162/necoa01160.

35. Marblestone AH, Wayne G, Kording KP. Toward an Integration of Deep Learning and Neuroscience. Frontiers in Computational Neuroscience. 2016;10. doi:10.3389/fncom.2016.00094.

36. Rombouts JO, Bohte SM, Roelfsema PR. How Attention Can Create Synaptic Tags for the Learning of Working Memories in Sequential Tasks. PLoS Comput Biol. 2015;11(3):e1004060. doi:10.1371/journal.pcbi.1004060.

37. Rombouts JO, Roelfsema P, Bohte SM. Neurally Plausible Reinforcement Learning of Working Memory Tasks. In: Advances in neural information processing systems; 2012. p. 1880–1888.

38. Rombouts JO, Bohte SM, Martinez-Trujillo J, Roelfsema PR. A learning rule that explains how rewards teach attention. Visual Cognition. 2015;23(1-2):179–205. doi:10.1080/13506285.2015.1010462.

39. Roelfsema PR, van Ooyen A, Watanabe T. Perceptual learning rules based on reinforcers and attention. Trends in Cognitive Sciences. 2010;14(2):64–71. doi:10.1016/j.tics.2009.11.005.

40. Roelfsema PR, Ooyen Av. Attention-Gated Reinforcement Learning of Internal Representations for Classification. Neural Computation. 2005;17(10):2176–2214. doi:10.1162/0899766054615699.

41. van Ooyen A, Roelfsema PR. A biologically plausible implementation of error-backpropagation for classification tasks. In: Supplementary Proceedings of the International Conference on Artificial Neural Networks; 2003. p. 442–4.

42. Roelfsema PR, Holtmaat A. Control of synaptic plasticity in deep cortical networks. Nature Reviews Neuroscience. 2018;19(3):166–180. doi:10.1038/nrn.2018.6.

43. Hochreiter S, Schmidhuber J. Long Short-Term Memory. Neural Computation. 1997;9(8):1735–1780. doi:10.1162/neco.1997.9.8.1735.

44. Gers FA, Schmidhuber J, Cummins F. Learning to forget: continual prediction with LSTM. 1999; p. 850–855. doi:10.1049/cp:19991218.

45. Gers FA, Schraudolph NN, Schmidhuber J. Learning precise timing with LSTM recurrent networks. Journal of machine learning research. 2002;3(Aug):115–143.

46. Monner D, Reggia JA. A generalized LSTM-like training algorithm for second-order recurrent neural networks. Neural Networks. 2012;25:70–83. doi:10.1016/j.neunet.2011.07.003.

47. Cho K, van Merrienboer B, Bahdanau D, Bengio Y. On the Properties of Neural Machine Translation: Encoder-Decoder Approaches. arXiv:14091259 [cs, stat]. 2014;.

48. Costa R, Assael IA, Shillingford B, de Freitas N, Vogels T. Cortical microcircuits as gated-recurrent neural networks. In: Guyon I, Luxburg UV, Bengio S, Wallach H, Fergus R, Vishwanathan S, et al., editors. Advances in Neural Information Processing Systems 30. Curran Associates, Inc.; 2017. p. 272–283.

49. Graves A, Schmidhuber J. Framewise phoneme classification with bidirectional LSTM and other neural network architectures. Neural Networks. 2005;18(5):602–610.

50. Graves A, Wayne G, Reynolds M, Harley T, Danihelka I, Grabska-Barwińska A, et al. Hybrid computing using a neural network with dynamic external memory. Nature. 2016;538(7626):471–476. doi:10.1038/nature20101.

51. Bakker B. Reinforcement learning with long short-term memory. In: Advances in neural information processing systems; 2002. p. 1475–1482.

52. Bakker B. Reinforcement learning by backpropagation through an LSTM model/critic. In: Approximate Dynamic Programming and Reinforcement Learning, 2007. ADPRL 2007. IEEE International Symposium on. IEEE; 2007. p. 127–134.

53. Hazy TE, Frank MJ, O’Reilly RC. Banishing the homunculus: Making working memory work. Neuroscience. 2006;139(1):105–118. doi:10.1016/j.neuroscience.2005.04.067.

54. Hazy TE, Frank MJ, O’Reilly RC. Towards an executive without a homunculus: computational models of the prefrontal cortex/basal ganglia system. Philosophical Transactions of the Royal Society B: Biological Sciences. 2007;362(1485):1601–1613. doi:10.1098/rstb.2007.2055.

55. Todd MT, Niv Y, Cohen JD. Learning to use working memory in partially observable environments through dopaminergic reinforcement. In: Advances in neural information processing systems; 2009. p. 1689–1696.

56. O’Reilly RC, Frank MJ, Hazy TE, Watz B. PVLV: the primary value and learned value Pavlovian learning algorithm. Behavioral neuroscience. 2007;121(1):31.

57. O’Reilly RC. The Leabra model of neural interactions and learning in the neocortex [PhD Thesis]. PhD thesis, Carnegie Mellon University, Pittsburgh, PA, USA; 1996.

58. O’Reilly RC. Biologically Plausible Error-Driven Learning Using Local Activation Differences: The Generalized Recirculation Algorithm. Neural Computation. 1996;8(5):895–938. doi:10.1162/neco.1996.8.5.895.

59. Everling S, Fischer B. The antisaccade: a review of basic research and clinical studies. Neuropsychologia. 1998;36(9):885–899.

60. Munoz DP, Everling S. Look away: the anti-saccade task and the voluntary control of eye movement. Nature Reviews Neuroscience. 2004;5(3):218–228. doi:10.1038/nrn1345.

61. Hallett PE. Primary and secondary saccades to goals defined by instructions. Vision research. 1978;18(10):1279–1296.

62. Brown MRG, Vilis T, Everling S. Frontoparietal Activation With Preparation for Antisaccades. Journal of Neurophysiology. 2007;98(3):1751–1762. doi:10.1152/jn.00460.2007.

63. Gottlieb J, Goldberg ME. Activity of neurons in the lateral intraparietal area of the monkey during an antisaccade task. Nature neuroscience. 1999;2(10):906.

64. Mello GBM, Soares S, Paton JJ. A Scalable Population Code for Time in the Striatum. Current Biology. 2015;25(9):1113–1122. doi:10.1016/j.cub.2015.02.036.

65. Tsao A, Sugar J, Lu L, Wang C, Knierim JJ, Moser MB, et al. Integrating time from experience in the lateral entorhinal cortex. Nature. 2018; p. 1. doi:10.1038/s41586-018-0459-6.

66. Zelinsky GJ. TAM: Explaining off-object fixations and central fixation tendencies as effects of population averaging during search. Visual cognition. 2012;20(4-5):515–545. doi:10.1080/13506285.2012.666577.

67. Choo FX, Eliasmith C. A spiking neuron model of serial-order recall. In: Proceedings of the 32nd Annual Conference of the Cognitive Science Society, ed. S. Ohlsson & R. Cattrambone; 2010. p. 218893.

68. Stewart TC, Tang Y, Eliasmith C. A biologically realistic cleanup memory: Autoassociation in spiking neurons. Cognitive Systems Research. 2011;12(2):84–92. doi:10.1016/j.cogsys.2010.06.006.

69. Zhang W, Luck SJ. Discrete fixed-resolution representations in visual working memory. Nature. 2008;453(7192):233–235. doi:10.1038/nature06860.

70. Cowan N. The Magical Mystery Four: How Is Working Memory Capacity Limited, and Why? Current Directions in Psychological Science. 2010;19(1):51–57. doi:10.1177/0963721409359277.

71. Bays PM, Husain M. Dynamic Shifts of Limited Working Memory Resources in Human Vision. Science. 2008;321(5890):851–854. doi:10.1126/science.1158023.

72. Van den Berg R, Awh E, Ma WJ. Factorial comparison of working memory models. Psychological review. 2014;121(1):124.

73. Ma WJ, Husain M, Bays PM. Changing concepts of working memory. Nature Neuroscience. 2014;17(3):347–356. doi:10.1038/nn.3655.

74. Brady TF, Alvarez GA. Hierarchical Encoding in Visual Working Memory: Ensemble Statistics Bias Memory for Individual Items. Psychological Science. 2011;22(3):384–392. doi:10.1177/0956797610397956.

75. Orhan AE, Sims CR, Jacobs RA, Knill DC. The Adaptive Nature of Visual Working Memory. Current Directions in Psychological Science. 2014;23(3):164–170. doi:10.1177/0963721414529144.

76. Barak O, Sussillo D, Romo R, Tsodyks M, Abbott LF. From fixed points to chaos: Three models of delayed discrimination. Progress in Neurobiology. 2013;103:214–222. doi:10.1016/j.pneurobio.2013.02.002.

77. Saxe A, Koh PW, Chen Z, Bhand M, Suresh B, Ng AY. On random weights and unsupervised feature learning. In: Proceedings of the 28th international conference on machine learning (ICML-11); 2011. p. 1089–1096.

78. Bouchacourt F, Buschman TJ. A Flexible Model of Working Memory. Neuron. 2019;103(1):147–160.e8. doi:10.1016/j.neuron.2019.04.020.

79. Mao T, Kusefoglu D, Hooks BM, Huber D, Petreanu L, Svoboda K. Long-Range Neuronal Circuits Underlying the Interaction between Sensory and Motor Cortex. Neuron. 2011;72(1):111–123. doi:10.1016/j.neuron.2011.07.029.

80. Aizawa H, Wurtz RH. Reversible inactivation of monkey superior colliculus. I. Curvature of saccadic trajectory. Journal of neurophysiology. 1998;79(4):2082–2096.

81. Bengio Y, Louradour J, Collobert R, Weston J. Curriculum learning. In: Proceedings of the 26th annual international conference on machine learning. ACM; 2009. p. 41–48.

82. Zaremba W, Sutskever I. Learning to execute. arXiv preprint arXiv:14104615. 2014;.

83. Martinolli M, Gerstner W, Gilra A. Multi-timescale memory dynamics in a reinforcement learning network with attention-gated memory. arXiv:171210062 [cs, q-bio, stat]. 2017;.

84. Rigotti M, Barak O, Warden MR, Wang XJ, Daw ND, Miller EK, et al. The importance of mixed selectivity in complex cognitive tasks. Nature. 2013;497(7451):585–590. doi:10.1038/nature12160.

85. Baddeley A. Working memory: looking back and looking forward. Nature Reviews Neuroscience. 2003;4(10):829. doi:10.1038/nrn1201.

86. Reggia JA, Monner D, Sylvester J. The computational explanatory gap. Journal of Consciousness Studies. 2014;21(9-10):153–178.

87. Sylvester JC, Reggia JA, Weems SA, Bunting MF. Controlling working memory with learned instructions. Neural Networks. 2013;41(Supplement C):23–38. doi:10.1016/j.neunet.2013.01.010.

88. Sylvester J, Reggia J. Engineering neural systems for high-level problem solving. Neural Networks. 2016;79(Supplement C):37–52. doi:10.1016/j.neunet.2016.03.006.

89. Eliasmith C. A Unified Approach to Building and Controlling Spiking Attractor Networks. Neural Computation. 2005;17(6):1276–1314. doi:10.1162/0899766053630332.

90. Berg Rvd, Ma WJ. A rational theory of the limitations of working memory and attention. bioRxiv. 2017; p. 151365. doi:10.1101/151365.

91. Vogel EK, Machizawa MG. Neural activity predicts individual differences in visual working memory capacity. Nature. 2004;428(6984):748–751. doi:10.1038/nature02447.

92. Oberauer K, Hein L. Attention to information in working memory. Current Directions in Psychological Science. 2012;21(3):164–169.

93. Sternberg S. High-Speed Scanning in Human Memory. Science. 1966;153(3736):652–654. doi:10.1126/science.153.3736.652.

94. Banks WP, Fariello GR. Memory load and latency in recognition of pictures. Memory & cognition. 1974;2(1):144–148.

95. Olivers CNL, Peters J, Houtkamp R, Roelfsema PR. Different states in visual working memory: when it guides attention and when it does not. Trends in Cognitive Sciences. 2011;15(7):327–334. doi:10.1016/j.tics.2011.05.004.

96. Wolfe JM. Saved by a Log: How Do Humans Perform Hybrid Visual and Memory Search? Psychological Science. 2012;23(7):698–703. doi:10.1177/0956797612443968.

97. Konecky RO, Smith MA, Olson CR. Monkey prefrontal neurons during Sternberg task performance: full contents of working memory or most recent item? Journal of Neurophysiology. 2017;117(6):2269–2281. doi:10.1152/jn.00541.2016.

98. Houtkamp R, Roelfsema PR. Matching of visual input to only one item at any one time. Psychological Research PRPF. 2009;73(3):317–326. doi:10.1007/s00426-008-0157-3.

99. Monsell S. Task switching. Trends in Cognitive Sciences. 2003;7(3):134–140. doi:10.1016/S1364-6613(03)00028-7.

100. Alport A, Styles EA, Hsieh S. 17 Shifting Intentional Set: Exploring the Dynamic Control of Tasks. 1994;.

101. Chatham CH, Frank MJ, Badre D. Corticostriatal Output Gating during Selection from Working Memory. Neuron. 2014;81(4):930–942. doi:10.1016/j.neuron.2014.01.002.

102. Myers NE, Rohenkohl G, Wyart V, Woolrich MW, Nobre AC, Stokes MG. Testing sensory evidence against mnemonic templates. eLife. 2015;4:e09000. doi:10.7554/eLife.09000.

103. Myers NE, Stokes MG, Nobre AC. Prioritizing Information during Working Memory: Beyond Sustained Internal Attention. Trends in Cognitive Sciences. 2017;21(6):449–461. doi:10.1016/j.tics.2017.03.010.

104. Rushworth MFS, Passingham RE, Nobre AC. Components of Switching Intentional Set. Journal of Cognitive Neuroscience. 2002;14(8):1139–1150. doi:10.1162/089892902760807159.

105. Houtkamp R, Roelfsema PR. The effect of items in working memory on the deployment of attention and the eyes during visual search. Journal of Experimental Psychology: Human Perception and Performance. 2006;32(2):423–442. doi:10.1037/0096-1523.32.2.423.

106. Soto D, Humphreys GW, Heinke D. Working memory can guide pop-out search. Vision Research. 2006;46(6):1010–1018. doi:10.1016/j.visres.2005.09.008.

107. Ort E, Fahrenfort JJ, Olivers CNL. Lack of Free Choice Reveals the Cost of Having to Search for More Than One Object. Psychological Science. 2017;28(8):1137–1147. doi:10.1177/0956797617705667.

108. de Vries IEJ, Van Driel J, Olivers CNL. Posterior *α* EEG Dynamics Dissociate Current from Future Goals in Working Memory-Guided Visual Search. Journal of Neuroscience. 2017;37(6):1591–1603. doi:10.1523/JNEUROSCI.2945-16.2016.

109. de Vries IEJ, Van Driel J, Karacaoglu M, Olivers CNL. Priority Switches in Visual Working Memory are Supported by Frontal Delta and Posterior Alpha Interactions. Cerebral Cortex. 2018;doi:10.1093/cercor/bhy223.

110. de Vries IEJ, Van Driel J, Olivers CNL. Decoding the status of working memory representations in preparation of visual selection. NeuroImage. 2019;191:549–559. doi:10.1016/j.neuroimage.2019.02.069.

111. Shadlen MN, Newsome WT. Neural basis of a perceptual decision in the parietal cortex (area LIP) of the rhesus monkey. Journal of neurophysiology. 2001;86(4):1916–1936.

112. Gold JI, Shadlen MN. The neural basis of decision making. Annual review of neuroscience. 2007;30.

113. Collins AGE, Frank MJ. Within-and across-trial dynamics of human EEG reveal cooperative interplay between reinforcement learning and working memory. Proceedings of the National Academy of Sciences. 2018;115(10):2502–2507. doi:10.1073/pnas.1720963115.

114. Masse NY, Yang GR, Song HF, Wang XJ, Freedman DJ. Circuit mechanisms for the maintenance and manipulation of information in working memory. Nature Neuroscience. 2019;22(7):1159–1167. doi:10.1038/s41593-019-0414-3.

